# Cerebrospinal Fluid Dynamics: Uncovering Alternative Blood Vessel Clearance Mechanisms

**DOI:** 10.1101/2024.06.13.598807

**Authors:** Qiuju Yuan, Senthil Kumaran Satyanarayanan, Suki Man-Yan Lee, Lingli Yan, Yaofeng Wang, Yan-Fang Xian, Liumin He, Yingying Zhou, Wutian Wu, You-Qiang Song, Huanxing Su, Zhi-Xiu Lin, Dajiang Qin

## Abstract

The pathways that run along the olfactory nerves crossing the cribriform plate and connecting to lymphatic vessels in the nasal cavity, have been identified as a crucial route for cerebrospinal fluid (CSF) outflow. However, the presence of a CSF efflux pathway through blood vessels in this region has yet to be clarified. This study aimed to elucidate the anatomical connections between the subarachnoid space and the bloodstream at the nasal epithelium and the venous drainage routes of the nasal epithelium in mice. Our findings demonstrated that CSF tracers could be drained not only through lymphatic vessels in the nasal cavity and cervical lymph nodes (CLNs), but also through the blood vessels in this area that extend to its venous drainage routes, including the facial and jugular veins. Additionally, we showed that ligation of CLNs neither impeded the influx and efflux of CSF tracers nor exacerbated Alzheimer’s disease (AD)-related pathology in AD mice. Our work reveals a previously unrecognized pathway for CSF drainage through blood vessels within the nasal mucosa. These findings provide insight into the efficient removal of waste products, facilitating optimal functioning of neural tissue within the susceptible tissue of our brains.

## Introduction

Fundamental to brain sustenance is the dynamic nature of its drainage systems, which have been a subject of research for over a century. Subarachnoid cerebrospinal fluid (CSF) surrounding the brain has long been considered to act as a sink for brain interstitial waste, including beta-amyloid, the culprit of Alzheimer’s disease (AD) (*1, 2*), although the mechanisms by which metabolites generated deep within the brain’s extracellular spaces are cleared into the subarachnoid CSF compartment is still subject to debate (*2, 3*). The mechanisms of CSF outflow are pivotal for maintaining intracranial pressure and central nervous system (CNS) homeostasis (*2*).

The current paradigm suggests a dual-outflow system for CSF to reach blood circulation. One pathway involves direct drainage of CSF into the venous blood through arachnoid projections, while the other pathway operates indirectly through the lymphatic system (*4*). The first view is based on work in the early 20^th^ century, now facing increasing challenges (*5, 6*). Only in more recent times has the significance of the lymphatic system in the drainage of CSF become evident (*7–12*). A compilation of studies has corroborated the drainage of CSF into nasopharyngeal lymphatics; however, despite imaging studies supporting these findings, the precise anatomical connections and their functional significance in CSF outflow remain elusive due to technical limitations (*10, 12–19*). Exploring the dynamics of CSF within the nasal mucosa in mice may provide a window into potential medical applications and a deeper understanding of neurological health. The utilization of tracers to map CSF pathways has revealed a significant outflow route through the sheaths encasing cranial nerves, particularly alongside the olfactory nerves traversing the cribriform plate. This porous bony structure demarcates the cranial from the nasal cavities, guiding CSF into the interstitial spaces of the nasal mucosa, where it can be absorbed by lymphatic vessels and subsequently channeled to cervical lymph nodes (CLNs) (*12*). This interstitial fluid (ISF), located in interstitial spaces, along with dissolved substances, bathes the cells and tissues, providing them with nutrients and oxygen (*20*). The contribution of blood vessels and lymphatic vessels to ISF absorption may vary depending on the local tissue conditions, including vascular densities and inflammation (*20*). However, it is generally agreed that, under normal circumstances, the absorption of interstitial fluid is predominantly accomplished by the blood vessels (blood capillaries) (*21–23*), while the lymphatic vessels help to remove any excess fluid that is not absorbed by the blood capillaries (*21–23*). Despite the evidence, the direct blood vessel contribution to CSF/ISF absorption has never been addressed in the nasal cavity. The implications of this anatomical journey are profound, especially considering the nasal cavity’s vast surface area, amplified by microvilli, and the rich vascularization beneath the epithelium coupled with a direct passage of venous blood into the systemic circulation. Thus, understanding the morphology and anatomy of the connection between the possible efflux pathway of CSF through blood vessels at this site, given the nasal mucosa, is crucial for identifying the etiopathogenesis of neurodegenerative diseases and associated diseases. This paper aims to broaden our understanding of traditional paradigms by providing compelling evidence derived from a study on CSF transport in mouse models.

Thus, in this investigation, we aimed to meticulously chart the anatomical pathways through which CSF communicates with the bloodstream within the nasal epithelium and its blood vessel drainage system in mice. Delving into the nuances of CSF dynamics, the study posits five objectives, including firstly to ascertain if CSF tracers can permeate into the lymphatic vessels located at the nasal epithelium and CLNs; secondly, to determine the possibility of CSF tracer drainage directly into the nasal mucosa’s blood vessels; thirdly, to explore the extension of tracer drainage into venous routes like the facial and jugular veins; fourthly, to evaluate the effects of ligating superficial cervical lymph nodes (sCLNs) and deep cervical lymph nodes (dCLNs) on the movement of CSF tracers; and finally, to discern whether this ligation influences AD pathology in transgenic mouse models. By investigating these pathways, the study seeks to unravel the complexities of CSF flow, potentially revolutionizing the understanding and treatment of neurodegenerative diseases. As will be elucidated in the subsequent sections, our findings depict CSF is drained directly into the bloodstream at the nasal mucosa, and this alternative pathway could compensate for the loss of cervical lymphatic system function in terms of CSF drainage. Strikingly, our data highlights that the conduits for CSF efflux flow are not primarily through the lymphatic vessels, as traditionally believed, but through an alternative blood vessel route.

## Materials and methods

### a) Animals

The study strictly followed the ethical guidelines and protocols for animal experiments established by The Chinese University of Hong Kong. The mice were kept in an environment with controlled temperature and humidity, adhering to a strict 12-hour light/dark cycle, and provided unlimited access to standard food and water. The research utilized specific strains: wild-type (WT) C57BL/6 mice and TgCRND8 (KM670/671NL + V717F) mice. TgCRND8 mice were bred in-house on a C57BL/6J background. The focus was on adult male mice, with each experimental group consisting of five mice two months old from the mentioned strains. A lymphatic reporter mouse line, Prox1-tdTomato, was also used to directly visualize the lymphatics in assessing the nature of the vessels in the nasal mucosa.

### b) Intra-cisterna magna (ICM) injection

The mice were sedated through an intraperitoneal (i.p.) administration of a combined solution of ketamine (100 mg/kg) and xylazine (10 mg/kg) dissolved in sterile saline and subsequently positioned on a heating pad (37 ± 0.5 °C) on a stereotactic apparatus followed by an incision on the scalp. Various tracer agents, including Evan Blue (EB) (10 µL, 5% in normal saline, 0.96 kDa, Sigma-Aldrich), Tetramethylrhodamine-conjugated dextran (10 µL, Tr-d10, 10 kDa, Thermo Fisher Scientific), the high-molecular-weight Alexa Fluor 555-conjugated Ovalbumin (10 µL, OVA555, 45 kDa, Thermo Fisher Scientific) or Alexa Fluor 488-conjugated ovalbumin (10 µL, OVA488, 45 kD, Thermo Fisher Scientific) and vehicle (artificial CSF) (10 µL), were directly injected into the cisterna magna at a controlled pace of 2 μL/min by employing a Hamilton syringe paired with a 33-gauge needle, operated with a syringe pump tailored to each experiment. The needle was kept intact for 1–2 minutes post-injection to prevent reflux.

### c) Tracer drainage in CNS-draining lymph nodes

To visualize the tracer drainage into dCLN and sCLN after the ICM injection (EB or Tr-d10) in a C57BL/6J mouse, an incision was made along the neck’s length above the clavicle, and sCLN was present in the upper part of the salivary gland. The sternocleidomastoid muscles were carefully moved aside, granting access to the dCLNs. Images of the tracer spread within the CNS-draining lymph nodes and associated tissues were acquired with a Zeiss microscope. After the injection procedure and tracer assessment, the mice were humanely euthanized using CO_2_.

### d) Lymph node surgical ligation and excision

The C57BL/6J and TgCRND8 mice were sedated as previously outlined and then securely positioned on cardboard in a supine position. The neck area was sanitized using iodine and 70% ethanol sequentially. Additionally, an ophthalmic solution was applied to the eyes to avert desiccation. For the sCLNs and dCLNs ligation process, a precise incision was executed midline, approximately 5 mm above the clavicle. The sCLNs were identified in the upper part of the salivary gland, while dCLNs were identified below the sternothyroid muscle. Subsequently, the afferent and efferent vessels of the sCLNs and the dCLNs were securely ligated employing a nylon suture (8–0 Nylon non-absorbable monofilament suture; Sigma Aldrich). The incision area was then sutured, followed by a subcutaneous antibiotics injection, and allowed to recover on a heating pad until gaining a response. After a stipulated period, tracers (EB or Tr-d10) were injected into cisterna magna as previously described to confirm the successful ligation and the rodents were humanely euthanized using CO_2_. Subsequently, these nodes and the whole brain were gently harvested, enabling a detailed immunohistochemical examination of the CNS-related lymph nodes, their affiliated vessels, and the entire brain for amyloid plaques. A parallel non-ligated group was used as a control.

### e) Tracer influx into the parenchyma of the brain

The entire brain sections of the ligated, non-ligated, and vehicle C57BL/6J mice were affixed to slides prepared with a gelatin coating, rinsed, and then sealed with a cover slip using a PBS/glycerol mixture for preservation. Imaging was performed on 6–8 coronal sections sequentially arranged from +0.5 mm to -2 mm relative to the bregma, utilizing a Zeiss digital microscope (Germany). This was done under a 5x objective lens, ensuring consistent exposure time, offset, and gain settings across all samples. The distribution of the tracers (Tr-d10 or OVA555) within these sections was meticulously quantified employing NIH Image J software for detailed analysis.

### f) Imaging and quantification of amyloid plaques

The brains of the TgCRND8 mice (ligated and non-ligated) were cut into 25 μm cryosections for staining of amyloid plaques. The brain slices were washed in PBS. The thioflavin S (TS) (Sigma-Aldrich, T1892, 1:5000) or Bam-10 antibody (Sigma-Aldrich, A5213, 1:1000) was employed to detect amyloid plaques and cerebral amyloid angiogenesis (CAA) in whole brain sections. In the process of TS staining, the brain sections undergoing free-floating preparation were subjected to multiple washes (3 times) with PBS. The sections were then treated with a TS working solution, prepared by diluting TS in a mixture of 0.5% in distilled water. Following a 1-minute incubation period, the sections were cleared of excessive dye using a TS washing solution of 70% ethanol in distilled water and rinsed thoroughly with PBS. Amyloid plaques and CAA in selective areas such as cortex, hippocampus, and thalamus were quantified using ImageJ software (ImageJ 1.39u, NIH, USA), with a systematic selection of cortical fields at 400× magnification for analysis. C57BL/6J mice were used as controls following the above procedure for quantifying amyloid-beta-associated gliosis.

### g) Tracer drainage in the nasal mucosa, facial and jugular veins

The C57BL/6J and Prox1-td Tomato mice were anesthetized, followed by the ICM injection with Tr-d10 or OVA55 or OVA488. The mice were then euthanized after 15, 30, 60, and 120 min without exsanguination and subsequently immersed in 4% paraformaldehyde (PFA) for 48 h fixation at room temperature.

To observe and dissect facial and jugular veins near the jaw’s hinge that extend towards the ear, the vascular area was exposed using fine surgical scissors. These vessels were carefully dissected for histological analysis.

A significant reference point for analyzing the nasal mucosa was the nasal dorsum of the mice, which displayed two distinct lines indicating the fusion between the maxillas on the sides and the ethmoid bone in the centre. By utilizing this anatomical feature as a guide, precise removal of the nasal bones and vomer was carried out, uncovering the septum’s dorsal and ventral boundaries. Subsequently, thorough sectioning of the palate was performed to detach the septum from the nasal wall on one side before repeating the process on the opposite side. Any remaining connections at the nasal tip were cut, and the mucosal tissues and the septum were entirely extracted. The nasal mucosa, encompassing the dorsal section and nasal septum, was carefully dissected and then immersed in 30 % sucrose at 4 °C overnight, setting the stage for OTC embedding and subsequent sectioning.

### h) Tracer drainage in the urinary bladder

Two hours after the ICM injection with Tr-d10 in ligated and non-ligated C57BL/6J, the mice were euthanized as described above. An incision was made along the midline of the lower abdomen, extending 1.5 cm above the pubic bone, revealing the bladder fully. The bladder was then gently retracted towards the head to uncover the bladder neck area. Employing a 22G needle, urine was collected, and its optical density was subsequently measured at 555 nm using a Spectrophotometer (Molecular Devices, USA).

### i) The levels of Aβ_40_ and Aβ_42_ in CSF

The TgCRND8 mice (ligated and non-ligated) under anaesthesia were positioned face down to expose their cisterna magna surgically. A laboratory-fashioned capillary tube with a tapered end was then used to penetrate the revealed meninges, allowing for the collection of CSF. The CSF samples were preserved at −80 °C for subsequent amyloid-beta (Aβ) analysis. Concentrations of Aβ_40_ and Aβ_42_ were quantified employing ELISA kits (#27713 and #27711; IBL, Fujioka, Gunma, Japan), strictly adhering to the guidelines provided by the manufacturer. Furthermore, the samples underwent normalization based on protein content using the BCA protein assay kit (#23225, Thermo Scientific Pierce, US) per the manufacturer’s instructions.

### j) Immunofluorescence staining

The frozen sections of the acquired samples, including brains, nasal mucosa, cervical lymph nodes, and blood vessel tissues, were rinsed with PBS for staining procedures. The following primary antibodies were used for immunostaining of mice tissues: goat anti-CD31 (1:500; R&D systems), goat anti-LYVE1 (1:1,000; R&D systems), rabbit anti-LYVE1 (1:1,000; AngioBio), rat anti-endomucin (1:1000, Santa Cruz Biotechnology), rabbit anti-αSMA (1:1000, Abcam), rabbit anti-haemoglobin (Sigma Aldrich), rabbit anti-IBA1 (1:1000, Wako), rat anti-GFAP (1:2000, Invitrogen), rabbit anti-neurofilament 200 (1:1000, Sigma Aldrich), Prussian blue (Sigma Aldrich), DAPI (1:10000, Abcam) and corresponding secondary antibodies (Alexa Fluor 488, 568 or 647, donkey anti-rabbit or rat or goat, 1:400) were used. The sections underwent a blocking process in a solution containing 0.3% Triton-X-100 with either 10% donkey serum or 1% BSA in 0.01M PBS for 90 min and incubation with specific primary antibodies overnight at 4°C. After washing in 0.01M PBS three times, sections were treated with corresponding Alexa Fluor® IgG (H+L) secondary antibodies (1:400) for an hour in darkness. After secondary antibody incubation, sections were washed with 0.01M PBS. For double immunostaining, the sections were incubated with specific primary antibodies at 4 °C overnight. After subsequent washes and secondary antibody incubation, sections were visualized under a Zeiss microscope. The labeled tissues/cells were quantified using ImageJ software (ImageJ 1.39u, National Institute of Health, USA), with a systematic selection of cortical fields at 400× magnification for analysis. The final average number of the three sections from each animal was used for analysis. Data collection and analysis were conducted blind to the experimental conditions.

## Results

### a) Dynamics of CSF tracer drainage within the nasal mucosa

The investigation into the drainage pathways of CSF tracers within the nasal region is pivotal for understanding the mechanisms of CSF clearance and its potential implications for neurological health. In this study, we first determined if the drainage of CSF tracer egresses from neural tissue into the nasal epithelium and subsequently enters the bloodstream within the nasal mucosa. Utilizing the tracer Tr-d10, we meticulously documented its presence across various components of the nasal mucosa, including the umbrella-like dorsal part and the nasal septum (**Fig 1a)**, highlighting the comprehensive nature of tracer dispersal within these regions (**Fig 1b**).

**Fig 1:**
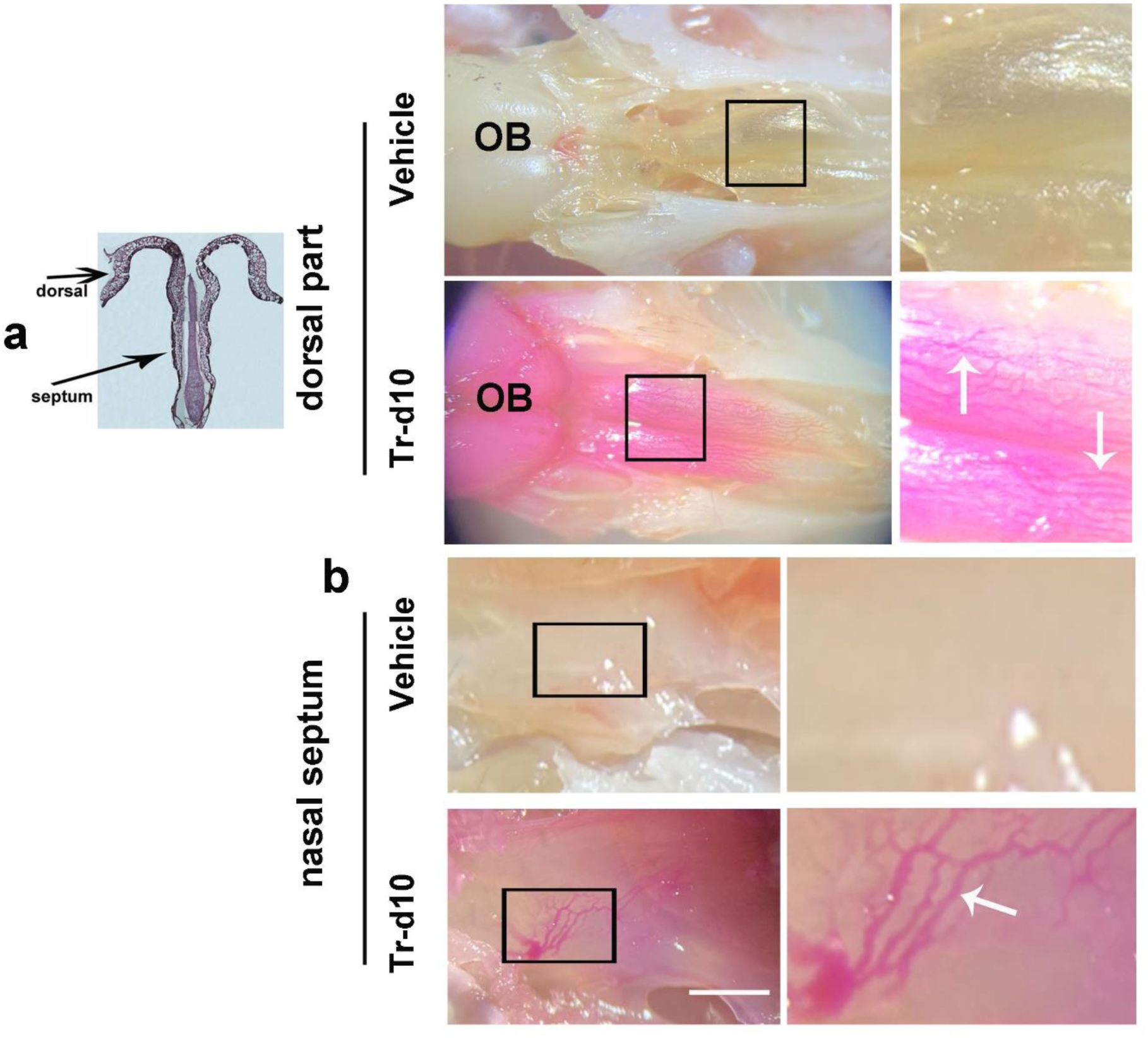
The CSF Tracer Distribution in Nasal Mucosa Regions. (a) Histological representation of the nasal mucosa, highlighting the dorsal part and the nasal septum. The arrows indicate the specific areas of interest for CSF tracer analysis. (b) Representative images of the nasal mucosa region showing the olfactory bulb (OB) and particular areas of interest, namely the umbrella-like dorsal part and the nasal septum in the mice. These images were captured 15 minutes after the ICM injection of either the vehicle or the Tr-d10 tracer. Widespread and extensive distribution of Tr-d10 was observed in the dorsal part and the septum of the nasal mucosa following Tr-d10 ICM injection. The magnified views on the right-hand side provide a closer examination of the tracer within the nasal vessels (arrows) in the areas enclosed by the black boxes. The scale bar represents 5 mm.

A pivotal aspect of this study involved the temporal analysis of Tr-d10 tracer dynamics within nasal mucosa post-ICM injection. Remarkably, tracer presence was noted in the lumen of blood vessels as early as 15 min following administration and was also distinct around 120 min (arrows in **Fig 2**), suggesting a rapid transit route from CSF spaces to vascular structures within the nasal mucosa (**Fig 2**). This observation underscores the efficiency of the nasal mucosa’s vascular network in facilitating tracer movement, potentially mirroring the pathways of CSF efflux.

**Fig 2:**
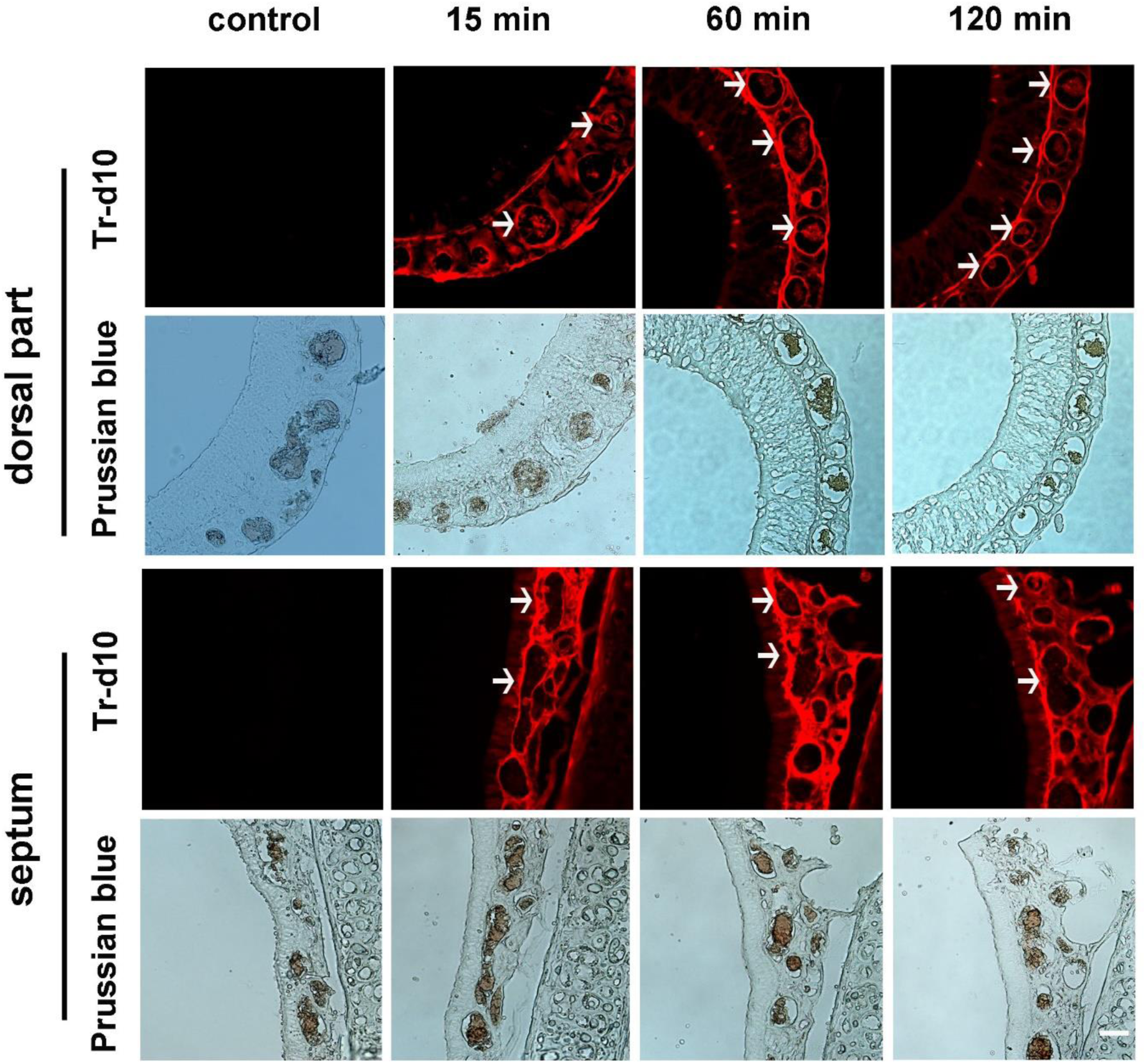
The CSF Tracer Distribution in the Blood Vessels of the Nasal Mucosa Post-ICM Injection. (a) Representative images of the dorsal part of the nasal mucosa at different time points post-ICM injection of Tr-d10 and vehicle control. At all the examined time points (15, 60, and 120 minutes) post-injection, the representative images demonstrate the presence of Tr-d10 within the vessel lumens (arrows), indicating rapid uptake of the CSF tracer. In contrast, the control images show no signal. The lower panel images are doubly labeled by Prussian blue staining of the same sections. This staining reveals the presence of iron content within the vessels, confirming that the vascular compartment serves as a conduit for the tracer distribution. (b) Representative images of the nasal septum of the nasal mucosa at different time points post-ICM injection of Tr-d10 and vehicle control. Similar to the dorsal part, the control images show no Tr-d10 signal. However, at 15-, 60-, and 120-min post-injection, the tracer is prominently visible within the vessel lumens (arrows). Additionally, these vessels are doubly labeled by Prussian blue staining, indicating the presence of iron content and confirming their nature as blood-filled vascular structures. The scale bar represents 20 µm.

Further investigations delved into the nature of tracer-filled vessels within the nasal mucosa, employing a combination of markers, including CD31, endomucin, hemoglobin, and Prussian blue, to characterize these conduits. Prussian blue, a unique stain, is instrumental in detecting the presence of ferric iron in tissues. Since ferric iron is a component of hemoglobin in the blood, the positive reaction to Prussian blue offers compelling evidence that the examined vessels contain blood. Crucially, these observations were made without exsanguination, meaning the blood within these vessels remained undisturbed, thus preserving their natural state. Endomucin, integral to the glycocalyx structure, stands out as a sialomucin with a substantial degree of O-glycosylation and a single transmembrane domain. This molecule is predominantly expressed in endothelial cells, with a specific prevalence in the endothelium of capillaries and veins, highlighting its vital role in vascular biology. Further, CD31, also known as platelet endothelial cell adhesion molecule (PECAM-1), is a protein often expressed on the surface of endothelial cells and is crucial in angiogenesis, a process of new blood vessel formation. The findings revealed that these tracer-bearing vessels were predominantly veins, a revelation that was consistently observed across various nasal mucosa regions, including the dorsal part (Tr-d10^+^/endomucin^+^/hemoglobin^+^/Prussian blue^+^) (**Fig 3a**) and septum (Tr-d10^+^/endomucin^+^/ Prussian blue^+^), in mice that had not undergone exsanguination (**Fig 3b**). This vein-centric tracer distribution pattern was further substantiated through the identification of vessels in prox1-Tom mice (endomucin^+^/ CD31^+^/Prussian blue^+^/hemoglobin^+^), reinforcing the venous predilection of CSF tracer drainage (**Fig 3c**). Meanwhile, the tracer was also taken up by lymphatic vessels in the nasal mucosa, as evidenced by the co-localization of Tr-d10 and lymphatic vessel endothelial hyaluronan receptor 1 (LYVE-1) (*24*)) (**Fig 3d**). The quantitative analysis of the vessels per section indicated a predominance of blood vessels over lymphatic vessels in tracer carriage (**Fig 3e**). This suggests that while lymphatic pathways may play a role in CSF drainage, the blood vessels within the nasal mucosa are the critical conduits for the efflux of CSF tracers. Such insights could have profound implications for our understanding of CSF dynamics, potentially influencing therapeutic strategies for diseases involving CSF circulation abnormalities. These observations collectively augment our comprehension of CSF clearance mechanisms, specifically the role of nasal mucosa as a conduit for CSF drainage into the blood vessel system. Identification of the veins as the primary vessels for tracer transport, as evidenced by specific marker co-localization, underscores the complexity of CSF dynamics.

**Fig 3:**
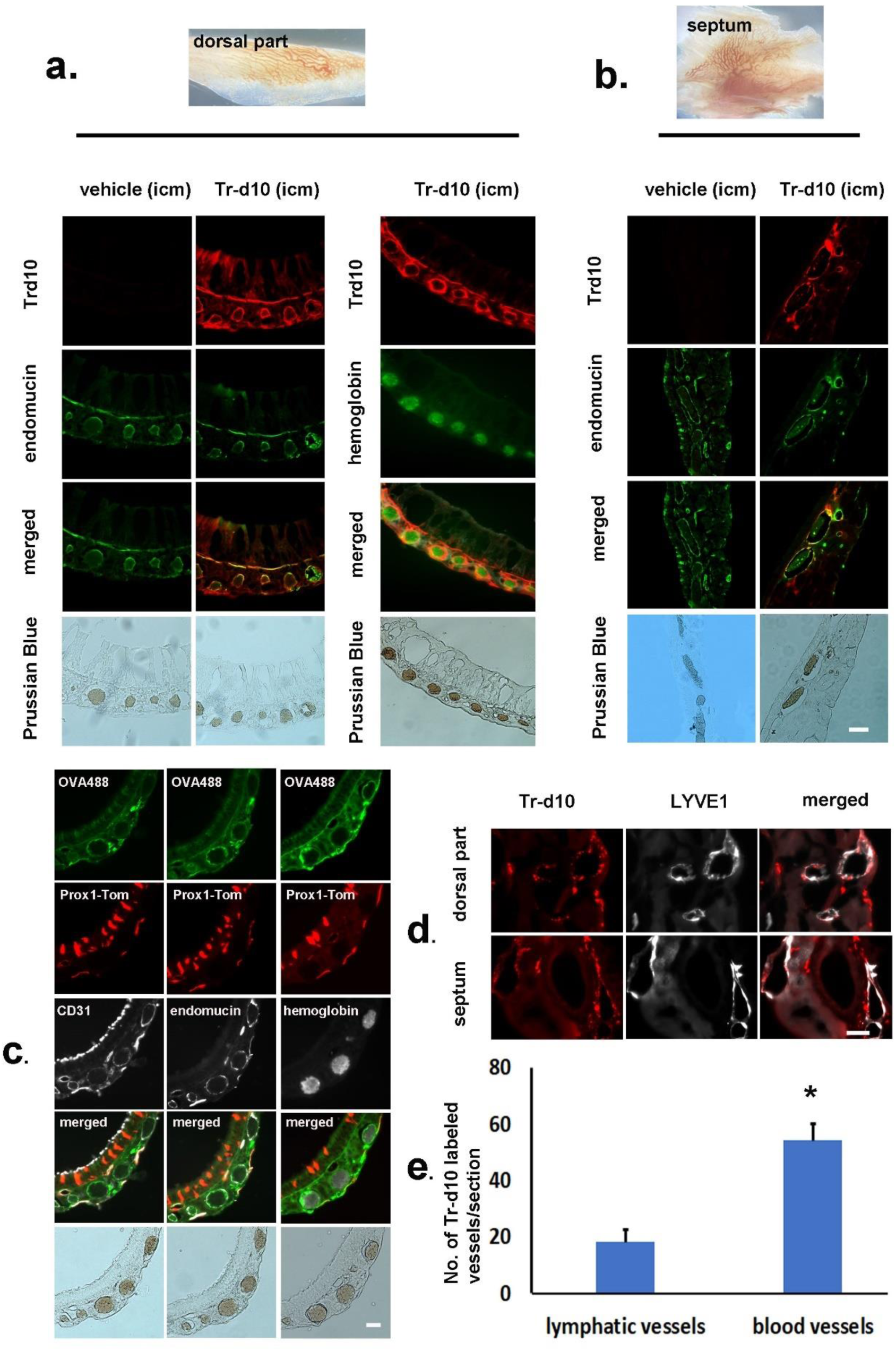
Characterization of Blood Vessels Containing the CSF Tracers in Nasal Mucosa regions. (a) Representative images of the dorsal part of the nasal mucosa. Two triple staining involving Tr-d10/endomucin/Prussian blue and Tr-d10/hemoglobin/Prussian blue are employed to identify the localization of Tr-d10 within the dorsal part of the nasal mucosa. The endomucin indicates endothelial cells of veins. The hemoglobin indicates the presence of red blood cells. Prussian blue staining reveals the presence of ferric iron. The co-localization of them characterized by Tr-d10+/endomucin+/Prussian blue+ or Tr-d10+/hemoglobin+/Prussian blue+ confirms the venous nature of Tr-d10 labeled vessels within the dorsal part of the nasal mucosa. (b) Representative images of the nasal septum. Similar to those in (a), a triple staining involving Tr-d10/endomucin/Prussian blue, was employed to illustrate the localization of Tr-d10 within the septum of the nasal mucosa. Their colocalization characterized by Tr-d10+/endomucin+/Prussian blue+ confirms the venous nature of Tr-d10 labeled vessels within the septum of the nasal mucosa. (c) Representative images of the dorsal part of the nasal mucosa in Prox1-Tom mice following ICM injection of a high-molecular-weight tracer, Alexa Fluor 488-conjugated Ovalbumin tracer (OVA488). Prox1 is one of the critical markers of lymphatic vessels. Three quadruple staining involving OVA488/Prox1-Tom/CD31/Prussian blue, OVA488/Prox1-Tom/endomucin/Prussian blue and OVA488/Prox1-Tom/hemoglobin/Prussian blue were employed to identify the localization of the CSF tracer OVA488 within the dorsal part of the nasal mucosa in Prox1-TdTom mice. The staining patterns characterized by OVA488+/Prox-1-Tom-/CD31+/Prussian blue+, OVA488+/Prox-1-Tom-/endomucin+/Prussian blue+, and OVA488+/Prox-1-Tom-/hemoglobin+/Prussian blue+ confirm their venous nature of OVA488 labeled vessels within the dorsal part of the nasal mucosa. (d) Representative images of the dorsal part and septum of the nasal mucosa in the wild-type mice following ICM injection of Tr-d10. Double staining involving Tr-d10/LYVE1 is employed to identify the localization of the CSF tracer Tr-d10. The LYVE1+ vessels indicated lymphatic vessels. Their colocalization characterized by Tr-d10+/LYVE1+ confirms the lymphatic nature of Tr-d10 labeled vessels within the nasal mucosa. (e) Quantification of the number of Tr-d10-bearing vessels per section of the nasal mucosa. The data highlight a higher number of blood vessels carrying the tracer compared to lymphatic vessels. The scale bar represents 20 µm.

### b) CSF Tracer Distribution in Lymph Nodes

The primary lymphatic drainage locations for the CNS have been identified as the dCLNs and sCLNs (*10, 17, 25*). Central to these observations, we examined the drainage and distribution of CSF tracers (EB and Tr-d10) within both dCLNs and sCLNs. In agreement with previous studies (*9, 10, 26–28*), we observed the EB and Tr-d10 tracer drainage in both dCLNs and sCLNs at 15 min onward following ICM tracers (**Fig 4a**). We further performed immunofluorescence staining in the sections of the CLNs using antibodies targeted against LYVE-1 to highlight the structure of the lymphatic nodes. As illustrated in **Fig.4b**, the paracortex and medulla of dCLNs and sCLNs simultaneously exhibited characteristic markers of lymphatic endothelial cells, namely LYVE-1, which were co-localized with Tr-d10. This finding highlighted the conventional lymphatic pathways for tracer flow in the dCLNs and sCLNs. The detection of CSF tracers in the dCLNs and sCLNs after ICM injection underscores their significance as a conduit, bridging the CNS and the lymphatic system. While the consistent observation of CSF tracers in dCLNs and sCLNs has established their involvement in CSF dynamics, we observed that the efferent vessel of sCLNs did not lead to dCLNs (**Fig 4c**). This awakens the quest to block both sCLNs and dCLNs if we aim to abolish the CSF outflow lymphatic pathway. Merely ligating the dCLNs may not completely block the cervical lymphatic pathway of CSF outflow.

**Fig 4:**
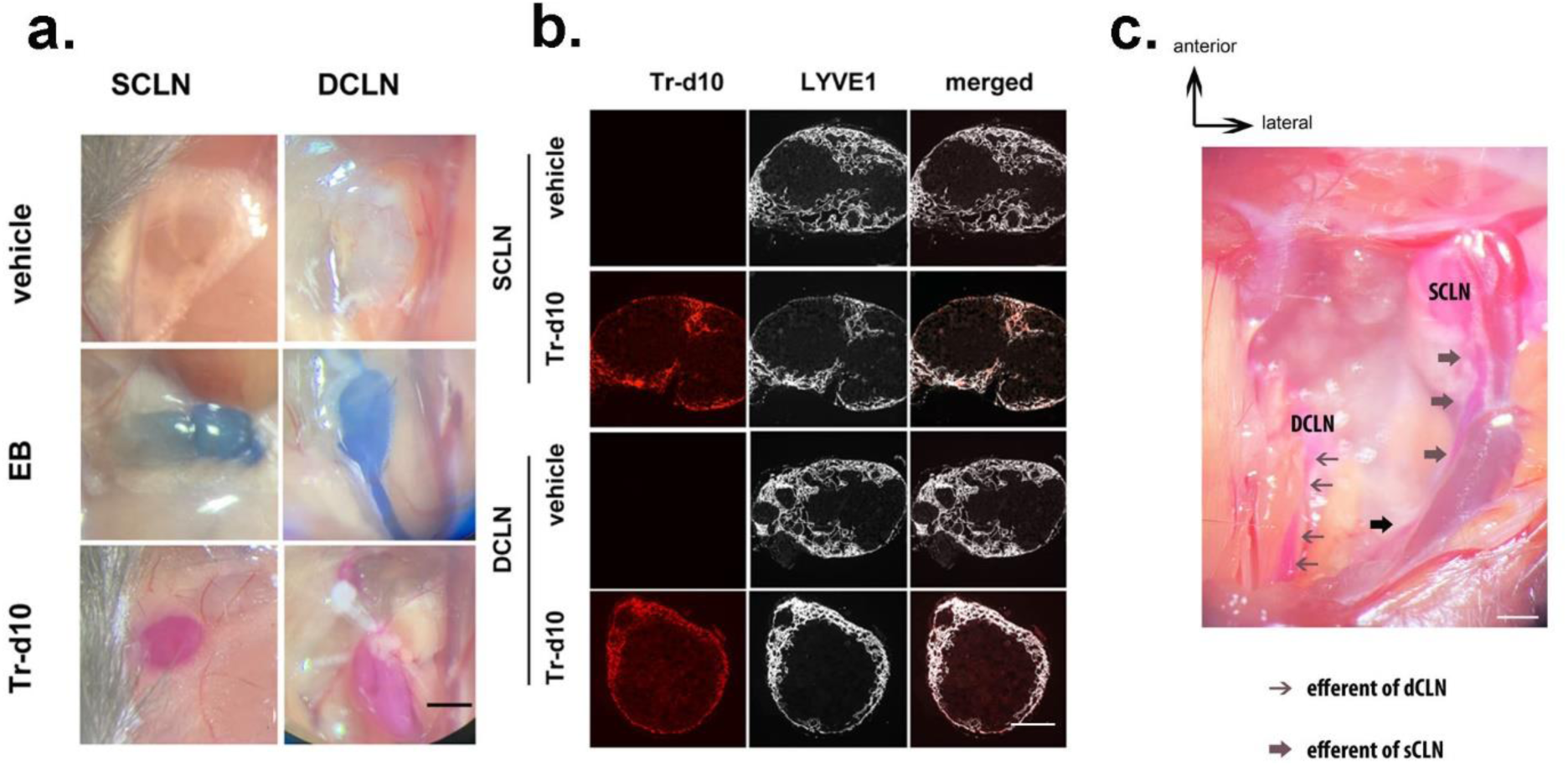
CSF tracers’ Drainage into Deep Cervical Lymph Nodes (dCLNs) and Superficial Cervical Lymph Nodes (sCLNs) (a) Representative images of dCLNs and sCLNs following ICM injection of tracers. The CSF tracers of EB and Tr-d10 were drained into dCLNs and sCLNs following the ICM injection. (b) Representative lymph node sections post-ICM Tr-d10 injection, displaying the presence of Tr-d10 in the LYVE1^+^ paracortex and medulla of sCLNs and dCLNs, evidencing the lymphatic drainage pathways. (c) Illustration of CSF tracer pathways in a dissected neck of the representative mice, highlighting the anterior-lateral orientation. Black arrows denote the efferent vessels of dCLNs, and arrowheads point to efferent vessels of sCLNs, showing distinct pathways without convergence post-ICM injection. The scale bar in (a) and (c) represents 1 mm. The scale bar in (b) represents 500 µm.

### c) Nasal epithelium’s venous drainage routes via facial and jugular veins

Exploring CSF tracer drainage into the nasal epithelium’s venous system, explicitly targeting its venous drainage pathways, including facial and jugular veins, represents a significant stride towards a better understanding CSF dynamic.

We subsequently delved into the mechanisms of CSF exit routes post ICM injection of Tr-d10 to map the drainage patterns and assess the involvement of various venous pathways in this process, including anterior facial vein (AFV), posterior facial vein (PFV), internal jugular vein (IJV) and external jugular vein (EJV). Following the administration of Tr-d10 *via* ICM injection, we observed the tracer’s presence in the lumen of AFV (**Figs 5a and 5b)** and PFV (**Figs 5a and 5c)** as early as 15 minutes post-injection, manifesting Tr-d10^+^ /Prussian blue^+^, persisting without exsanguination. This rapid appearance underscores the efficiency of facial veins in facilitating CSF drainage. The peak of Tr-d10 coverage within the facial veins at the 30-minute mark, rather than at 120 minutes (**Fig 5d)**, highlights a swift drainage process, which was further affirmed by the co-localization of Tr-d10 with endomucin and Prussian blue (**Fig 5e)**, marking the endothelial cells and providing visual confirmation of the tracer within these veins.

**Fig 5:**
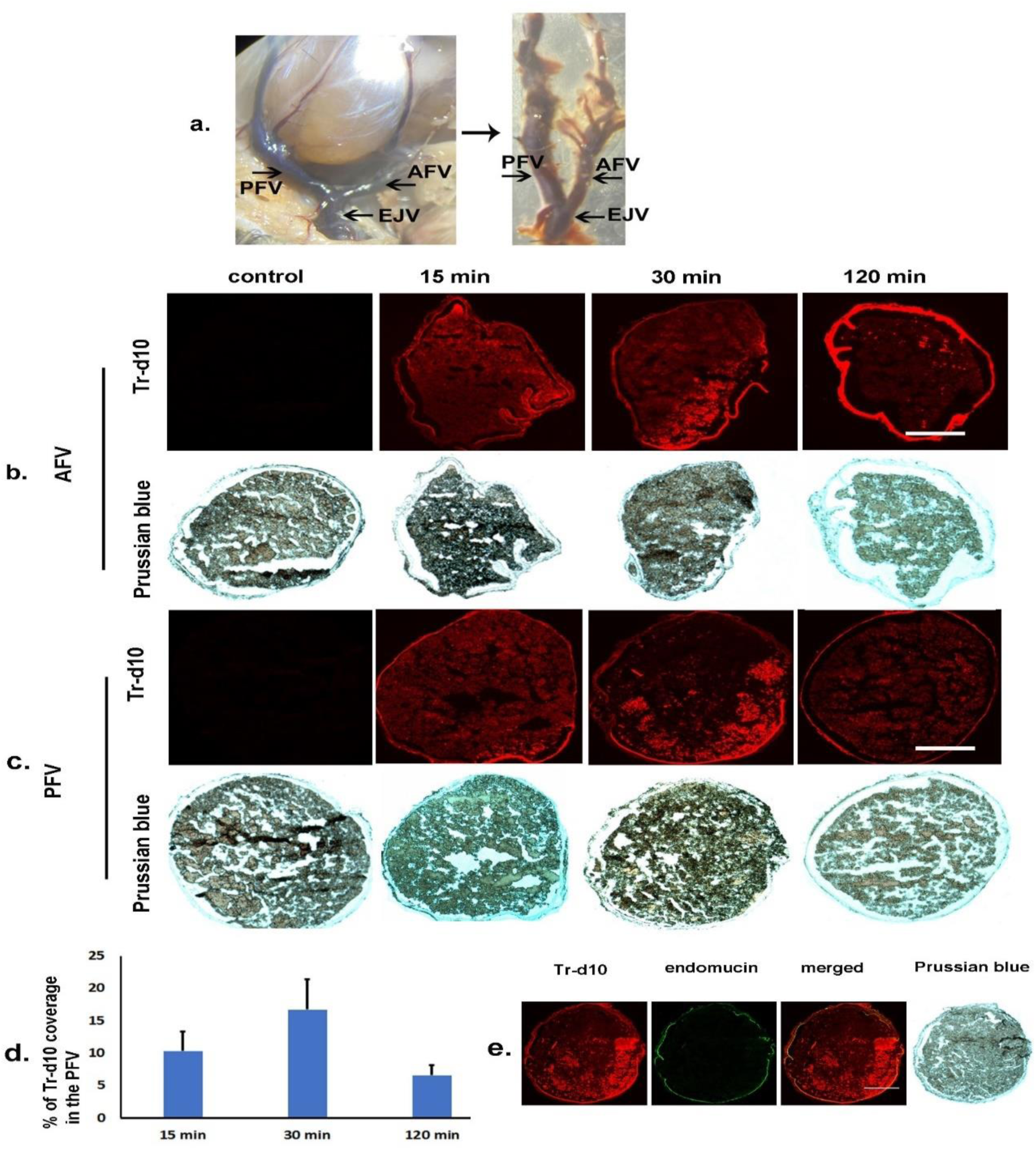
The CSF Tracer Drainage Through Facial Vein Pathways After ICM Injection. (a) Macroscopic images of the head and neck vasculature of the mice identifying the anterior facial vein (AFV), posterior facial vein (PFV), and external jugular vein (EJV), providing anatomical context for the drainage study. (b) Representative images of the AFV following ICM injection of Tr-d10 at various time points (control, 15, 30, and 120 minutes) showing the presence of the tracer within the vein. The lower panel shows corresponding Prussian blue staining of the same sections, indicating the presence of blood cells within the vascular structures. (c) Similar to (b) showing representative images and Prussian blue staining of the PFV at the same time points post-ICM injection, indicating the drainage of Tr-d10 through the PFV. (d) Quantification of Tr-d10 area fraction within the PFV over time (mean ± SEM), demonstrating a peak in tracer presence at 30 minutes and a decrease at 120 minutes, suggesting a rapid CSF clearance through this pathway. (e) A triple staining involving Tr-d10/endomucin/Prussian blue employed to validate the venous specificity of the Tr-d10-bearing vessel. Their colocalization, characterized by Tr-d10+/endomucin+/Prussian blue+, confirms the venous specificity of CSF drainage. The scale bar represents 250 µm.

Expectedly, the EJV was identified as another critical exit route for CSF, with a pattern similar to the facial vein as revealed by the Tr-d10 tracer (**Fig 6a and 6b)**, delineating the venous specificity (Tr-d10^+^/endomucin^+^/ Prussian blue^+^) of CSF drainage (**Fig 6c**).

**Fig 6:**
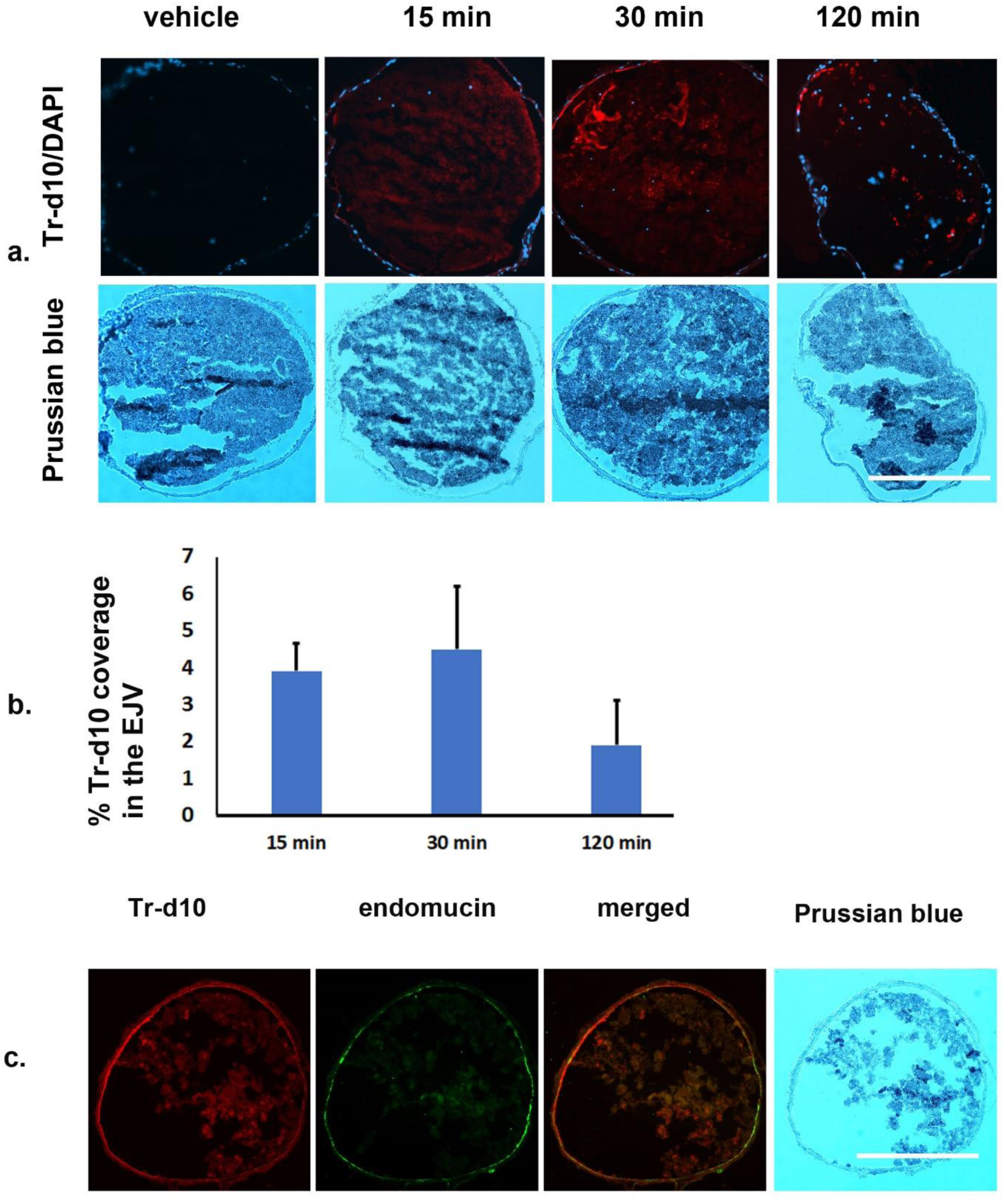
CSF Tracer Drainage Through the EJV Pathway After ICM Injection. (a) Representative images displaying the presence of Tr-d10 in the EJV at 15, 30, and 120 minutes after ICM injection compared to the vehicle. The bottom panels show the corresponding Prussian blue staining of the same sections, which marks the presence of iron, indicative of blood cells. (b) Quantification of Tr-d10 area fraction within the EJV over time (mean ± SEM). The data shows tracer presence within the EJV, with peak coverage at 30 minutes and a decrease at 120 minutes, suggesting active CSF drainage through the vein. (c) A triple staining involving Tr-d10/endomucin//Prussian blue in the EJV employed to validate the venous specificity of the Tr-d10-bearing vessel. Their colocalization, characterized by Tr-d10+/endomucin+/Prussian blue+, confirms the venous specificity of CSF drainage through the EJV. The scale bar represents 600 µm.

Additionally, the IJV, the critical venous drainage route of the nasal cavity (**Fig 7a)**, was also identified as another critical exit route for CSF, with Tr-d10 detected within as early as 15 min following ICM of Tr-d10 (stars in **Fig 7b and 7c)**, contrasting sharply with the absence of significant Tr-d10 levels in the common carotid artery with Tr-d10^-^ /DAPI^+^/Prussian blue^+^ in Fig 7b (an arrow **in Fig 7b)** and Tr-d10^-^/αSMA ^+^/LYVE-1^-^/CD31^+^/Prussian blue^+^ in Fig 7c (arrows in **Fig 7c),** indicating the venous specificity (Tr-d10^+^/αSMA^+^/LYVE-1^-^/CD31^+^/hemoglobin^+^/Prussian blue^+^) of CSF drainage.

**Fig 7:**
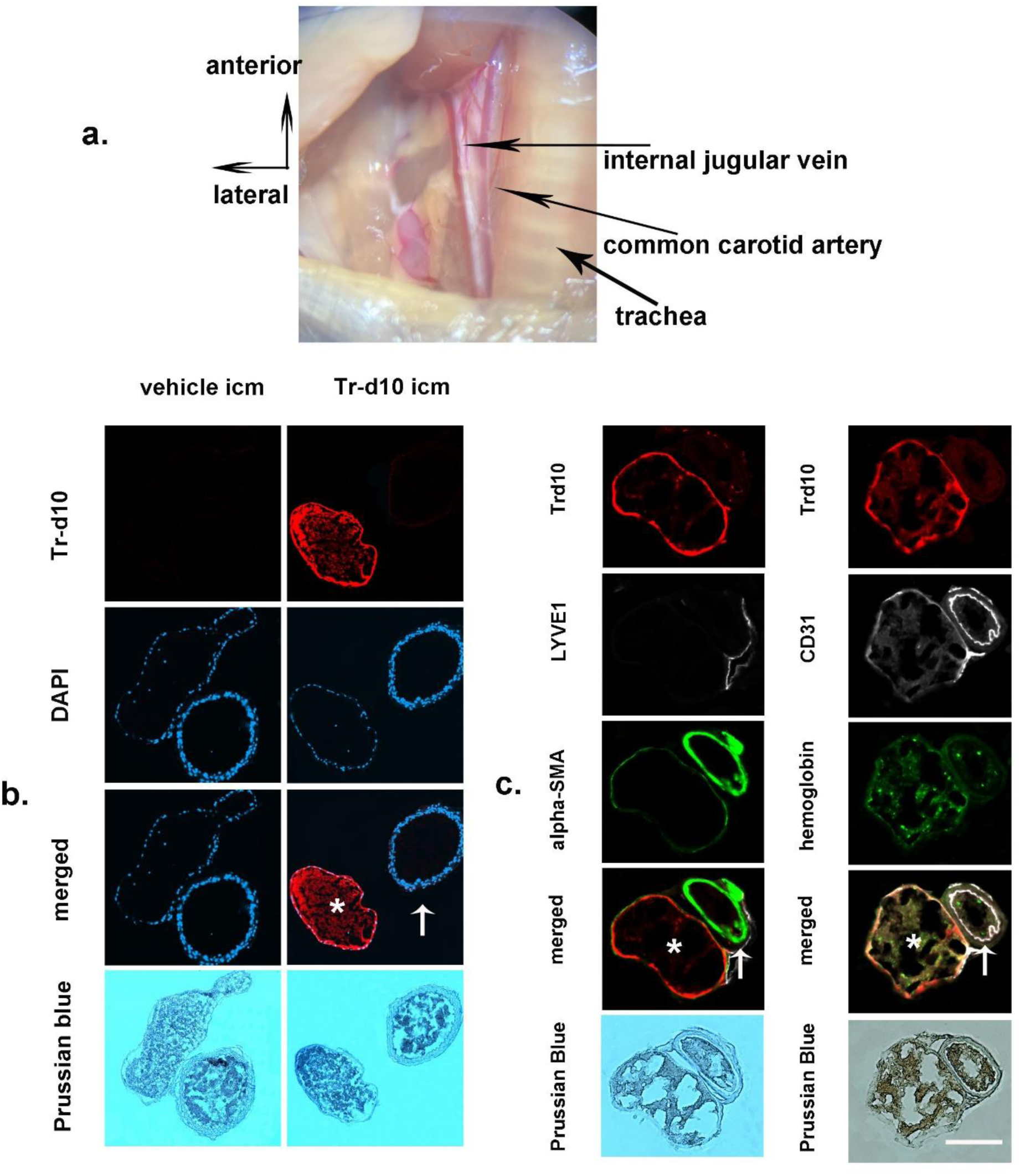
CSF Tracer Drainage through the IJV pathway After ICM Injection. (a) The neck region’s anatomical orientation depicted in an anterior-lateral view, highlighting critical structures such as the internal jugular vein (IJV), common carotid artery, and trachea. The IJV is recognized as a crucial venous drainage pathway for the nasal cavity. (b) Representative images illustrating the IJV and common carotid artery by nuclei staining with DAPI after injecting Tr-d10 and the vehicle. The vehicle control image does not show any Tr-d10 signal. However, at 15 minutes post-injection, the tracer is visible within the IJV’s lumen (a star). The IJV is distinguished by a thinner wall, as revealed by DAPI staining. In contrast, the adjacent carotid artery, which has a thicker wall (an arrow), does not display the presence of the CSF tracer. (c) A quadruple staining involving Tr-d10/αSMA/LYVE1/Prussian blue employed to identify the localization of Tr-d10 within the tissues. The staining indicates that Tr-d10 is in the IJV (a star) characterized by αSMA+/LYVE-1-/Prussian blue+ staining, which presents a thinner wall. In contrast, Tr-d10 is not located in the carotid artery, demonstrating αSMA+/LYVE-1-/Prussian blue+ staining, but has a thicker wall (an arrow). In an adjacent section, a quadruple staining using Tr-d10/CD31/hemoglobin/Prussian blue further confirms the localization of Tr-d10 in the CD31+/hemoglobin+/Prussian blue+ IJV, while the adjacent carotid artery does not show Tr-d10 localization. The scale bar represents 200 µm.

### d) Post-Ligation Changes in Lymphatic Nodes and Brain Parenchyma

Following the ligation of dCLNs, particularly when combined with ICM tracers’ injection, we observed pronounced alterations in the afferent vessel’s structure (**Fig 8a)**. These changes were evidenced by significant enlargement of the afferent vessel with the accumulation of visible tracers (EB and Tr-d10) (arrows in **Fig 8a)** and further absence of tracers within the node **(Fig 8a).** This suggests an impairment in the extracranial lymphatic drainage system and a cessation of tracer dye flowing to these lymph nodes, an effect which becomes more pronounced after the ligation of the dCLNs (*9, 28*). Further, when we analyzed the changes in the nodes with the ligated group, histological sections confirmed that the ligated nodes displayed markers Tr-d10^-^/ LYVE-1^+^ compared with the non-ligated Tr-d10^+^/ LYVE-1^+^, suggesting that the ligation prevented the entry of tracers into both dCLNs and sCLNs (**Fig 8b**).

**Fig 8:**
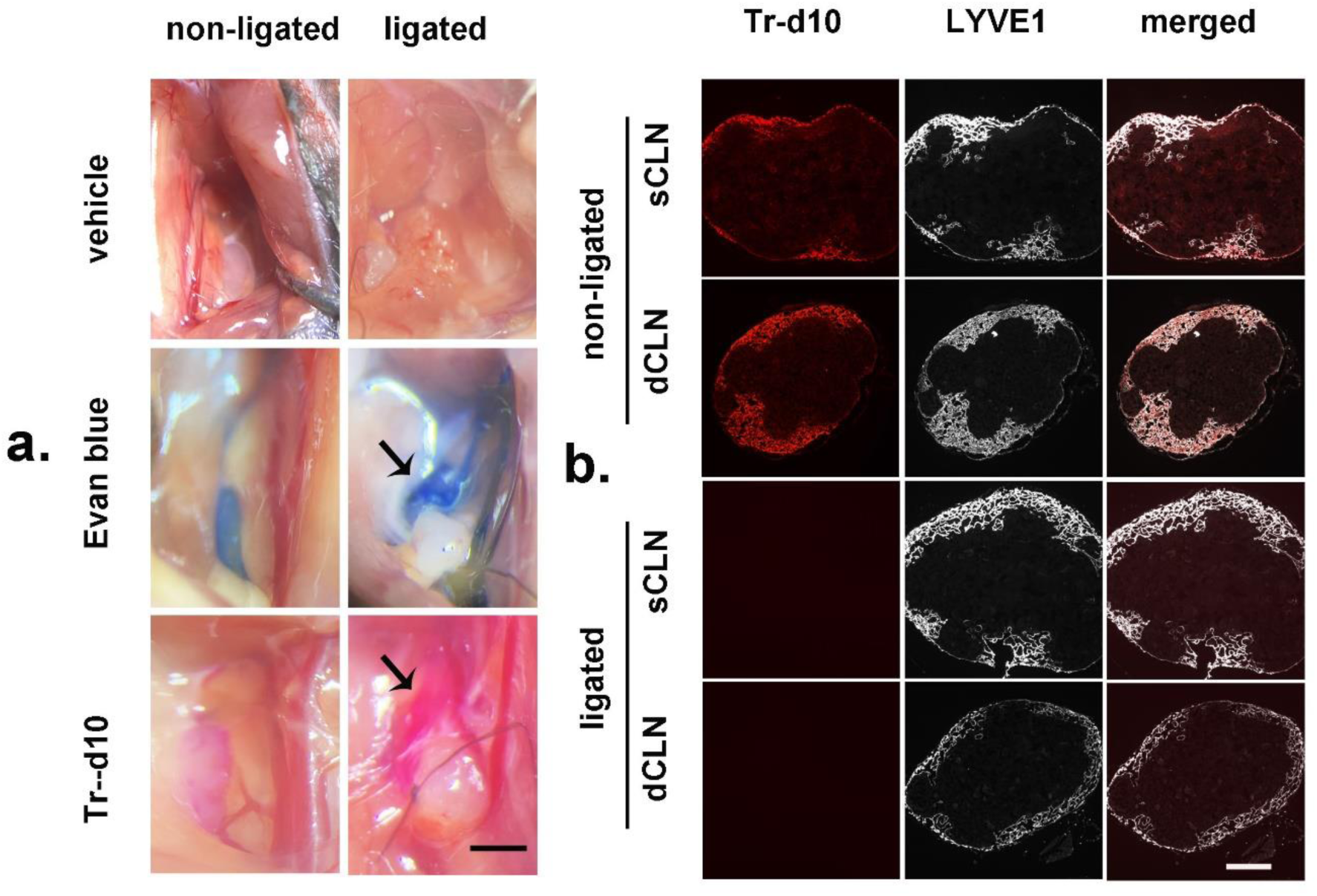
CLNs Ligation Prevents the Entry of the Tracer into CLNs. (a) A comparison of cervical lymph node regions between those that were ligated and those that were not ligated after ICM injection of EB and Tr-d10. The ICM injection of EB and Tr-d10 resulted in these tracers in the dCLNs and their afferent and efferent vessels. When the dCLNs were ligated, the afferent vessels exhibited enlargement with accumulated CSF tracers (EB and Tr-d10) observed at 15 minutes after the injection (arrows), in contrast to the afferent vessels of non-ligated nodes, indicating a successful interruption of CFS tracer entering the CLNs by the ligation. Additionally, the ligation caused the absence of visible EB or Tr-d10 in the dCLNs. (b) Representative lymph node sections stained with LYVE1 immunostaining after Tr-d10 ICM injection. Non-ligated dCLNs displayed the presence of Tr-d10 following ICM injection, whereas ligated nodes showed no Tr-d10 signal, indicating that ligation prevented the entry of the tracer. The scale bar in (a) represents 1mm. The scale bar in (b) represents 150 µm.

Further, our objective was to discern the impact of cervical node ligation (dCLNs and sCLNs) on the flow of CSF into the brain’s parenchyma and the clearance of CSF tracers from the brain. Through histological examination and quantitative analyses measuring tracer accumulation within the brain’s parenchyma, it was found that ligation of the CLNs did not significantly alter the influx of Tr-d10 into the brain’s parenchyma within 1 h post-ICM injection (**Figs 9a and 9c)**. Similarly, the efflux of this tracer from the brain, assessed at 72 h post-injection, was comparable between ligated and non-ligated models (**Figs 9a and 9c)**, indicating complete drainage of the tracer from the brain. This pattern of unaffected tracer dynamics was consistent with that of another tracer, OVA555, under the same experimental conditions (**Figs 9b and 9d**).

**Fig 9:**
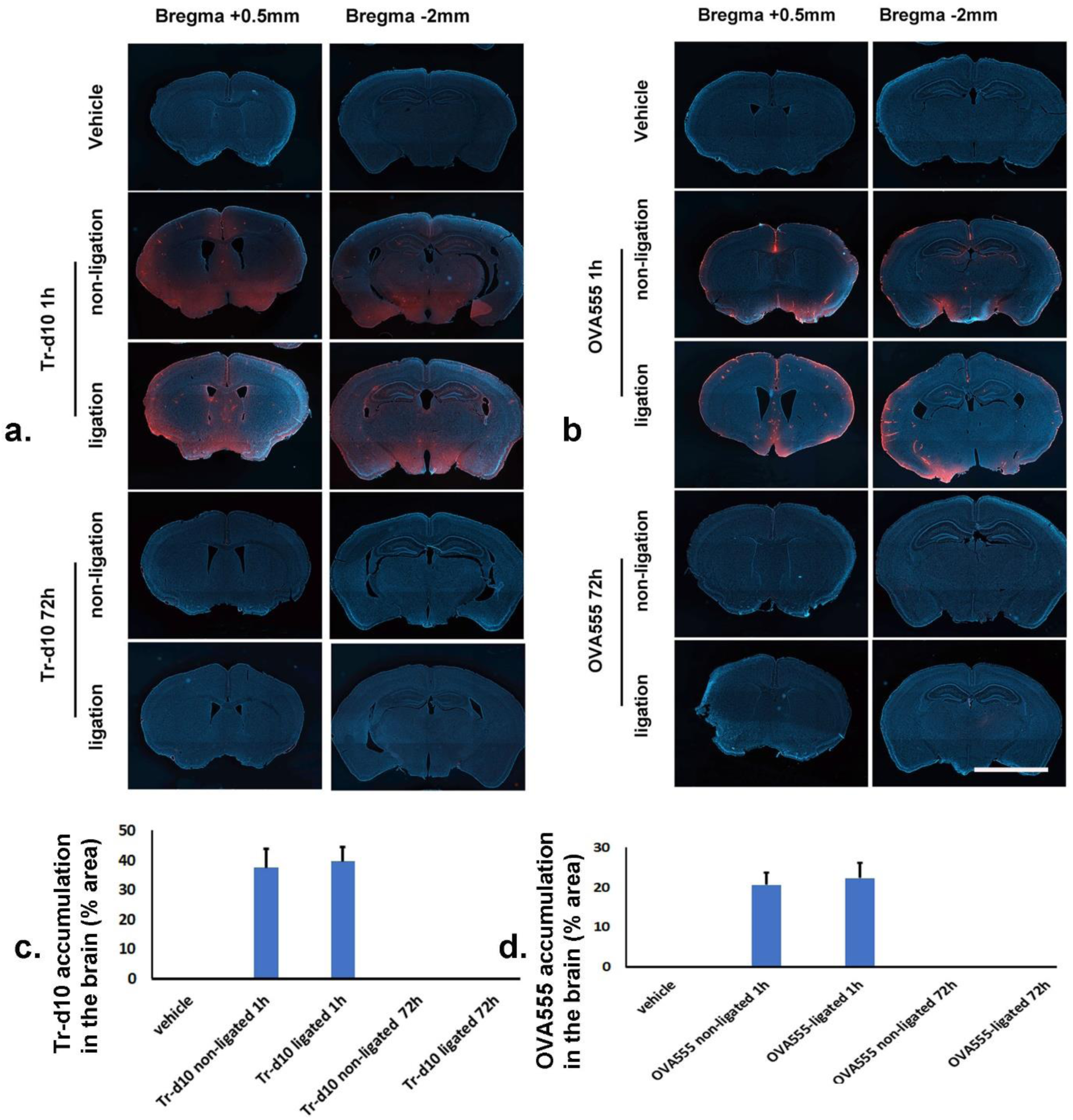
CLNs Ligation Does Not Affect Brain CSF Influx and Efflux. (a) Representative brain sections showing DAPI and Tr-d10 at 1 h and 72 h post-injection following either CLNs non-ligation or ligation. (b) Representative brain sections showing DAPI and OVA555 at 1 h and 72 h post-injection following either CLNs non-ligation or ligation. Representative images at 1h post-injection display the presence of Tr-d10 or OVA555 in the brain in either the non-ligated or ligated group, indicating the ligation could not inhibit CSF tracer influx. Similarly, the absence of Tr-d10 or OVA555 in the brain in either the non-ligated or ligated group at 72 h post-injection indicates the ligation could not inhibit CSF tracer clearance (c) Quantification of Tr-d10 area fraction. (d) Quantification of OVA555 area fraction. Data in (c) and (d) is presented as mean ± SEM. (a–d) show no significant differences between non-ligated and ligated groups.

Furthermore, the drainage of Tr-d10 into the urine of ligated mice remained unaltered 2 h after ICM tracer injection (**Fig 10**). These findings suggest that cervical lymph node ligation does not significantly impede CSF from entry into the brain’s parenchyma or its subsequent clearance, indicating the existence of alternative or compensatory pathways for CSF circulation in the presence of cervical lymphatic system disruption.

**Fig 10:**
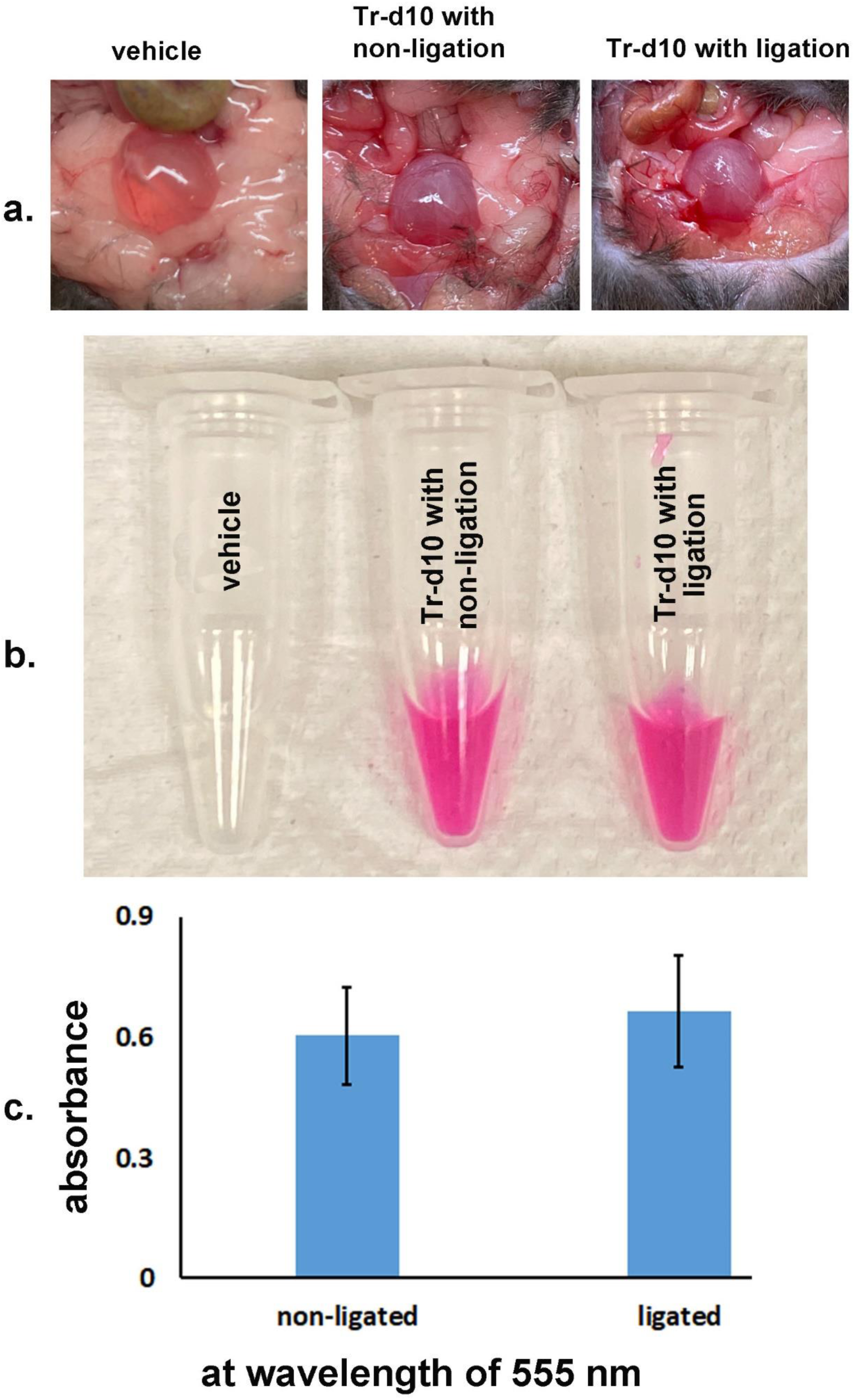
CSF Tracer Clearance into Urine Despite CLNs Ligation. (a) Representative dissected lower abdomen showing urinary bladder at two h following ICM injection of vehicle (left), Tr-d10 with non-ligation (center), and Tr-d10 with ligation (right) of CLNs. (b) Visual representation of collected urine samples from the vehicle (left), non-ligated (center), and ligated (right) mice post-ICM injection of vehicle and Tr-d10. The presence of pink coloration indicates the tracer’s passage into the urine. (c) Quantitative analysis of Tr-d10 in urine samples based on absorbance at a wavelength of 555 nm. The graph depicts similar absorbance levels for non-ligated and ligated groups, suggesting unaffected tracer elimination from the CNS into the urine.

Next, we wanted to know whether the clearance rate of CSF tracers of a similar scale is attributed to compensatory drainage through blood vessels, even in cases where the cervical lymphatic drainage pathway is completely obstructed. We have indeed observed an increasing trend of CSF tracer in the PFV (posterior facial vein) after ligating lymph nodes (**Fig 11**).

**Fig 11:**
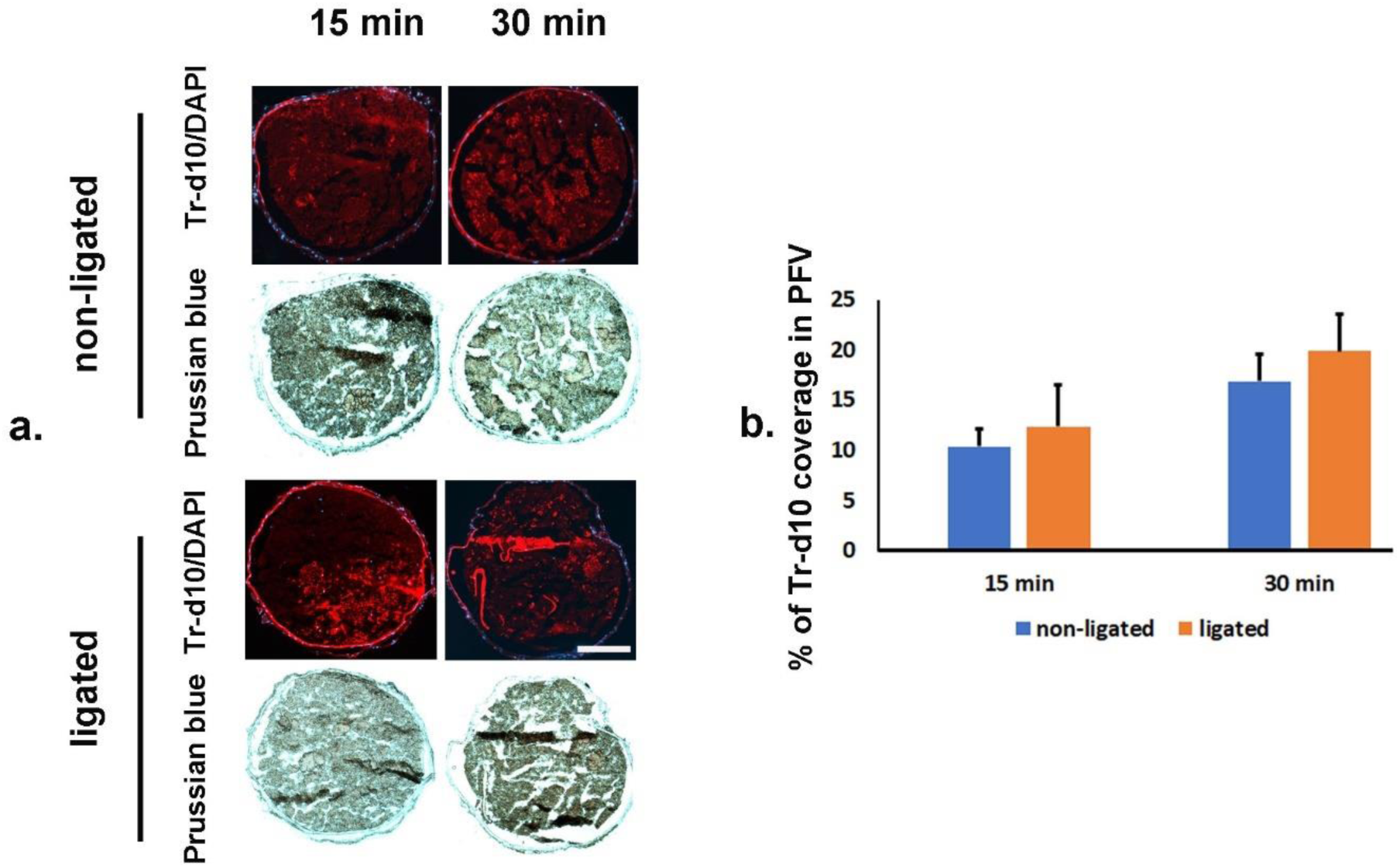
CSF Tracer Clearance via the Posterior Facial Vein (PFV) Following CLNs Ligation. (a) Representative images of PFV sections from mice subjected to CLNs ligation or non-ligation conditions at 15 and 30 min after ICM injection of Tr-d10. The upper panel shows the presence of Tr-d10 in PFV in non-ligated mice at both time points. In contrast, the lower panel indicates the distribution of Tr-d10 in PFV after CLNs ligation, illustrating the impact of ligation on CSF tracer clearance patterns. (b) Quantitative analysis of the percentage of Tr-d10 coverage in PFV, comparing non-ligation and ligation conditions at 15 and 30 minutes post-ICM injection. The graph demonstrates an increasing trend in the proportion of tracer Tr-d10 within PFV over time, with a more pronounced increase observed in the ligated group, suggesting compensatory drainage through blood vessels when the cervical lymphatic pathway is obstructed. The scale bar represents 250 µm.

### e) Cervical node ligation did not aggravate Aβ pathology, and Aβ associated glial reactions

We further aimed to discern the impact of cervical node ligation on Aβ pathology and associated glial reactions in TgCRND8 mice, a transgenic model of AD. A meticulous quantitative analysis conducted a month post-ligation revealed no significant increase in CSF Aβ levels (Aβ_40_ and Aβ_42_) in ligated subjects compared to non-ligated controls (**Fig 12a**). Moreover, the extent of Aβ pathology, encompassing plaques and cerebral amyloid angiopathy (**Figs 12b, 12c, and 12d**), did not show exacerbation within critical brain regions such as the cortex, hippocampus, and thalamus in ligated mice. Similarly, the investigation into Aβ-associated gliosis, marked by the presence of microglia (**Fig 12e)** and astrocytes (**Fig 12f)**, demonstrated no significant aggravation in the ligation group versus the non-ligation group (**Fig 12g**).

**Fig 12:**
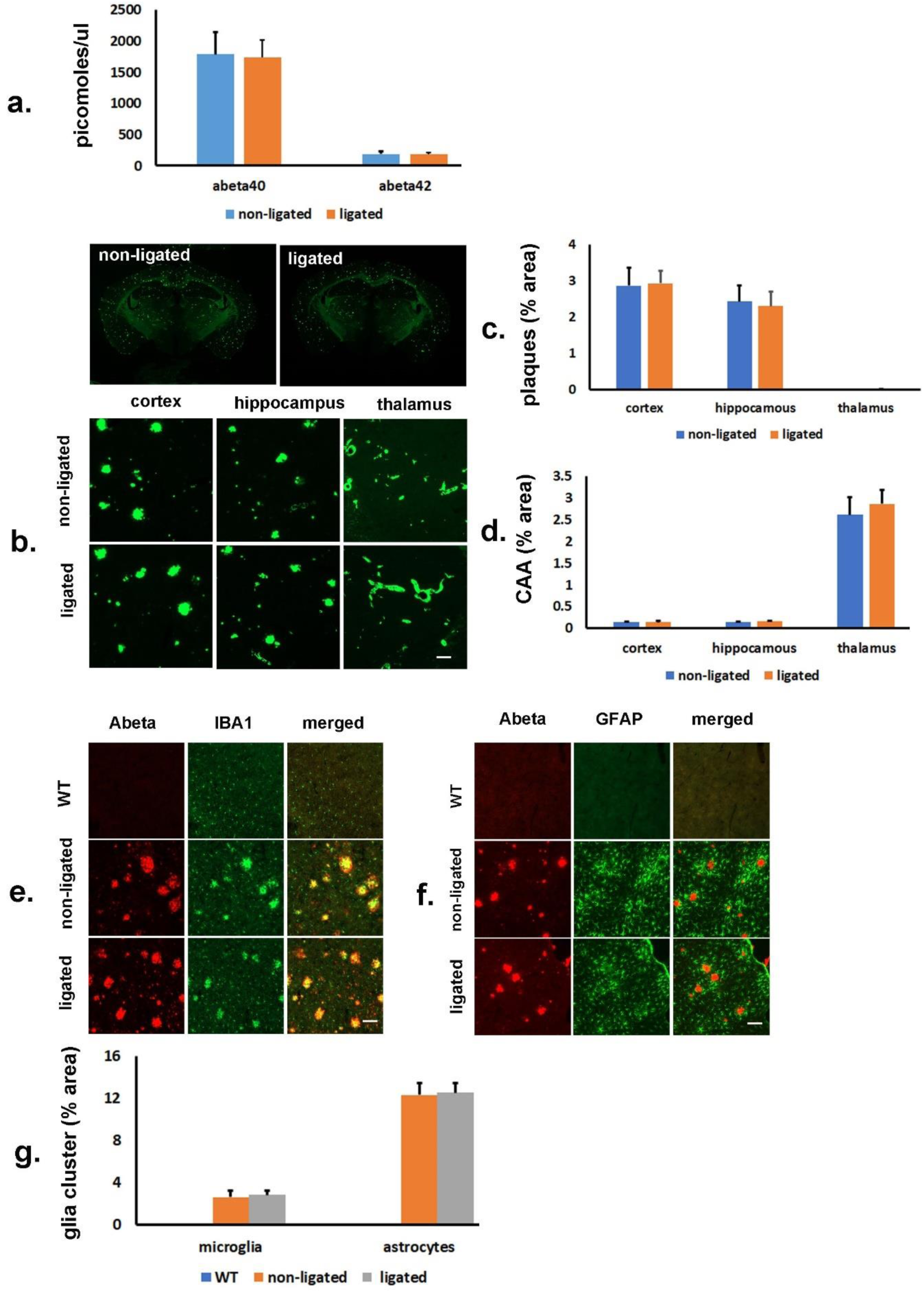
CLNs Ligation Does Not Affect Aβ Pathology and Glial Responses in TgCRND8 AD Mice. (a) Bar graph depicting CSF levels of Aβ_40_ and Aβ_42_ peptides in transgenic TgCRND8 mice with and without cervical lymph node ligation. Data indicate no significant change in the concentration of these peptides between the non-ligated and ligated groups one month post-ligation. Each bar represents the mean ± SEM of the levels of Aβ_40_ and Aβ_42_ in the CSF measured using ELISA. (b) Young-adult TgCRND8 mice at the age of two months were submitted to CLNs ligation or control procedures. Aβ pathology, including Aβ plaques and CAA in the cortex, the hippocampus, and the thalamus, was assessed seven months after the ligation as revealed by thioflavin S (TS) staining. (c-d), Quantification of Aβ pathology (c) Aβ plaques, (d) CAA in the cortex, the hippocampus, and the thalamus of TgCRND8 mice from the two groups. Each bar represents the mean ± SEM of coverage of Aβ plaques or CAA in the three regions of the brain. The graph shows no notable difference between non-ligated and ligated mice in CAA and Aβ pathology. (e-f) Representative images of (e) IBA^+^ microglia and (f) GFAP^+^ astrocytes clustering around Aβ plaques in the cortex of TgCRND8 mice from the two groups. (g) Quantification of glia clustering, including microglia and astrocytes clustering around Aβ plaques in the cortex of TgCRND8 mice from the two groups. Each bar represents the mean ± SEM of coverage of microglia or astrocytes clustering in the context of the brain. The graph shows no significant difference in glial activation between the ligated group and non-ligated controls. The scale bar represents 20 µm.

## Discussion

This study aimed to delineate the CSF drainage mechanisms in mice and revealed multifaceted pathways for CSF clearance. In this study, we have anatomically characterized the CSF outflow pathways in the olfactory region of mice by employing CSF tracer infusions. We then confirmed that alternative blood vessel drainage at the nasal mucosa might compensate for impaired lymphatic drainage, indicative of alternative CSF clearance pathways beyond the lymphatic system. Our research provides a comprehensive chain of evidence, including the drainage of CSF tracer into the nasal mucosa, widespread dispersion of this tracer within the nasal mucosa, and its uptake by blood vessels of the nasal mucosa. Furthermore, the tracer enters the nasal mucosa’s venous drainage pathways: the facial and jugular veins. These findings indicate that blood vessels participate in the absorption and drainage of CSF in the nasal mucosa, allowing its entry into the bloodstream. Through experiments involving bilateral sCLN and dCLN ligation, we further demonstrate that blood vessel drainage of CSF can compensate for the lymphatic system’s CSF drainage function. Moreover, we observed that ligation of cervical lymph nodes in AD mice does not exacerbate amyloid-beta and its associated pathological changes. This study presents the inaugural and significant finding that CSF can be directly assimilated into the bloodstream *via* the dense blood vessels of the nasal mucosa, indicating a highly effective new pathway for CSF drainage. This efficiency stems from the extensive capillary network and large surface area for absorption and transportation into the bloodstream in the nasal mucosa. Compared to the lymphatic system, the blood vessels provide significantly greater blood flow (over 100 times more) (*29*). This accelerated blood flow enables swift elimination of CSF and the brain’s metabolic waste, including amyloid-beta, ensuring a favourable brain environment.

The discharge of CSF through the cribriform plate into the nasal cavity represents the first described pathway for CSF outflow (*4*). Furthermore, numerous studies across a diverse range of species have corroborated that tracers introduced into the CSF are indeed expelled through the cribriform plate, reinforcing this pathway’s significance in CSF outflow (*10, 12, 30–41*). It was not until more recently, however, that the importance of the lymphatic system to CSF drainage became apparent (*7–12, 17*). A compilation of studies has corroborated the drainage of CSF into the cervical lymph nodes (*4, 10, 12, 19, 42*). However, the anatomical mechanisms of this CSF drainage pathway remain elusive (*4*). Three main possible anatomical pathways have been suggested (*4*). The initial pathway involves perineural routes that travel alongside the olfactory nerve sheaths, traversing the interstitial tissue between them to reach the lymphatic vessels in the submucosa. This theory is referred to as the “open cuff model,” in which it is postulated that the perineural sheath cells vanish beyond the cribriform plate, creating a space for cerebrospinal fluid (CSF) to flow into the interstitial area outside the skull. Within this interstitial space, the CSF is absorbed by the initial lymphatics in the olfactory and respiratory submucosa. The second pathway entails perineural routes along the olfactory nerve sheaths, directly connecting to the lymphatic vessels in the submucosa. This concept is known as the “closed cuff model,” which proposes that the perineural space forms a widened pouch or cul-de-sac, allowing the lymphatic vessels to merge with the perineural cells and gain direct access to CSF flowing along the olfactory nerve. The third pathway suggests lymphatic vessels crossing the plate might connect directly to the subarachnoid space (SAS). In both the second and third models, it is plausible that CSF tracers enter and fill the lymphatics as a path of least resistance rather than spreading through the interstitial spaces of the submucosa. However, more recent data show a massive, dispersed distribution of CSF tracers in the nasal mucosa (*17, 19*). In line with the previous studies, we observed massive coloration of Tr-d10 in the nasal mucosa. The fluorescence extended to olfactory epithelium and submucosa. The findings suggest a preference for the first model, as it uniquely enables CSF to propagate via the interstitial space of the submucosa. This is not surprising given that the endothelial layer of the olfactory nerve sheath becomes increasingly thinner in the nasal mucosa, which allows CSF and solutes to access the interstitial spaces easily (*4*).

Our hypothesis is based on the first model, which posits that if CSF can be discharged into the interstitial space of the nasal mucosa and subsequently taken up by lymphatic vessels, absorption by capillaries and even veins should also be feasible. It draws on established principles regarding the differential roles of blood and lymphatic vessels in ISF absorption. Blood and lymphatic vessels play crucial roles in fluid circulation and regulation within the human body. While blood vessels absorb tissue fluid *via* capillary walls, lymphatic vessels take up tissue fluid through the lymphatic system. Under typical conditions, blood vessels account for a substantial proportion of tissue fluid absorption, approximately 90%, as capillaries feature walls are exceptionally permeable, enabling the transfer of solutes and fluids directly into the bloodstream. The vascular system propels this process, leveraging the pressure difference between arteries and veins to facilitate fluid absorption and circulation. Conversely, lymphatic vessels account for a modest portion of tissue fluid absorption, approximately 10%. Operating alongside the vascular system, they collect tissue fluid and waste, channeling these through lymph fluid to lymph nodes and subsequently into the bloodstream. The efficiency of lymphatic vessels in fluid absorption is limited, predominantly depending on the pressure of tissue fluid and peristaltic movement within the vessels to propel lymph movement (*22, 23*).

The nasal mucosa stands out for its dense vascular network, providing a notably expansive area suitable for absorption (*43*). Additionally, studies in humans and other mammals have revealed the presence of fenestrated endothelial walls in capillaries and veins situated beneath and directly adjacent to the basal cell layers of the nasal epithelium (*44–46*). The unique characteristics of blood vessels within the nasal mucosa, characterized by their heightened permeability, facilitate the swift and efficient systemic absorption of substances (*43*). This underpins the potential for intranasal drug delivery to be as effective as intravenous administration (*47, 48*). Given that medications can penetrate the bloodstream after crossing the nasal epithelium’s basal cell layer barrier, it stands to reason that CSF in the interstitial space similarly enters the bloodstream. In our research, we distinguished vessel types using markers like CD31, endomucin, and alpha-SMA, which were also found to be present in collecting lymphatic vessels (*49*). A definitive feature sets it apart from blood because lymphatic fluid’s lack of red blood cells renders it colorless. Therefore, our study focused on tracking the CSF tracer in mouse models without exsanguination. Firstly, we can employ methods to identify red blood cells with Prussian blue staining and employing immunohistochemistry with anti-hemoglobin antibody to analyze blood vessel characteristics. The second, which is more important, is more conducive to ascertaining whether the CSF tracer was within vessel lumens rather than adhering to vessel exteriors, a scenario possibly influenced by artificial factors like lymphatic leakage. Additionally, immunohistochemical staining with anti-endomucin antibody helped us assess the venous attributes of blood vessels. The lumen diameter served as another differentiator, especially for facial and jugular veins in mice, which can exceed 300 µm, contrasting with the much narrower diameters of larger collecting lymphatic vessels, typically around 20-30 µm.

In our study, we observed the presence of CSF tracer in the veins of the olfactory mucosa. This could be attributed to absorption through the capillaries of the olfactory mucosa or direct absorption through the veins in that area, aligning with previous studies that have shown that some veins beneath the olfactory epithelium exhibit a porous structure similar to capillaries (*44–46*). Our observations confirmed the entry of the CSF tracer into the bloodstream *via* blood vessels in the nasal mucosa, ensuring its inevitable passage into the olfactory mucosa’s downstream venous drainage. This venous drainage from the nasal cavity comprises a sophisticated web of veins differing in size and location (*4, 50*).

The major venous drainage routes of the nasal cavity include the facial vein and pterygoid venous plexus. The pterygoid venous plexus ultimately drains into the maxillary vein and retromandibular vein. The retromandibular vein splits into two segments: the anterior branch merges with the facial vein to create the common facial vein, which subsequently empties into the IJV. Meanwhile, the posterior branch combines with the posterior auricular vein to establish the EJV. In short, both IJV and EJV participate in nasal cavity venous drainage, although there are differences in the facial vein drainage pathway between humans and rodents (4, 50). In rodents, facial veins drain into the EJV. Instead, facial veins drain into the IJV (4, 50). Our research uncovered that the CSF tracers were channeled into the nasal mucosal venous drainage pathways, including the AFV, PFV, IJV, and EJV. These results differ from the findings of another research group that has made significant discoveries in the field of CSF drainage (10). This contrasts with Ma et al. (10) and other scholars in the CSF drainage research domain (12), who, despite identifying facial veins, did not report CSF tracer presence within these veins. We speculate that this variation could stem from the different observational techniques applied. While those groups utilized in vivo imaging methods offering a comprehensive perspective, such approaches may overlook weaker signals, limiting their detection of CSF tracer in certain vessels and primarily observing its presence in lymphatic channels (4, 10). Our in vivo study confirmed that the CSF tracer was detected exclusively in lymphatic vessels and CLNs of prox1 tdTom mice. With this method, we were also unable to observe CSF tracer in the facial and jugular veins (**Suppl. 1**). This may be due to the lower concentration of tracer in the veins, resulting in insufficient signal strength to be detected by this method. The concentration of CSF tracer within the lumen certainly affects the signal strength. We did observe weaker signals when we injected the tracer with lower concentrations into CSF (data not shown). It is worth noting that the diameter of the facial and jugular veins (300-1200 µm) is substantially larger than cervical collecting lymphatic vessels (20-30 µm). Thus, the cross-sectional area of these veins is over one hundred to even thousands of times larger compared to the lymphatic vessels, suggesting that any CSF tracer entering the bloodstream through nasal mucosa vessels would be considerably diluted in these veins. This dilution likely contributes to the challenges in detecting the tracer in veins with in vivo imaging.

In our study, we observed the presence of CSF tracer in the veins of the nasal mucosa and the downstream pathways of the facial and jugular veins. A pivotal question arises: does the CSF tracer directly enter these blood vessels, or is it an indirect process? Common consensus holds that CSF permeates the nasal mucosa via the cribriform plate, gets absorbed by lymphatics, and drains into CLNs before entering the systemic circulation. Is it possible that the CSF tracer observed in the mentioned veins is a CSF tracer that has been drained via lymphatics back into the systemic circulation and subsequently reaches the blood vessels of the whole body? To confirm or rule out this possibility, we explored the presence of CSF tracer in veins and arteries elsewhere, including the femoral vein and artery, without detecting significant tracer presence at any experimental time points (**Suppl. 2**), nor within the carotid artery close to the IJV. Crucially, even after ligating the sCLNs and dCLNs to block lymphatic CSF drainage, the CSF tracer still appeared in the veins and urine. These findings strongly indicate that the CSF tracer in these blood vessels is not indirectly introduced through the systemic circulation but instead directly absorbed, leading to detectable concentrations.

Beyond the primary CSF drainage route via the nasal mucosa, earlier research has identified two supplementary pathways: through the spinal nerve roots and towards the sacral lymph nodes (SLN) (10, 51). To determine if these pathways contribute to compensatory drainage following lymphatic obstruction, we investigated the presence of CSF tracers at these locations. We detected CSF tracers moving within the spinal subarachnoid space, heading towards the lower extremity of the spine (**Suppl. 3a and 3b**). However, our results indicate that spinal nerves may not significantly contribute to CSF drainage following either ICM EB or Tr-d10, as evidenced by the minimal drainage of the CSF tracer Tr-d10 through spinal nerves in both ligated and non-ligated models (**Suppl. 3a, 3b and 3c**). Immunofluorescence analysis revealed the presence of neurofilament 200 (NF200), a neurofilament marker indicating nerve fibres within the brachial and lumbar plexus, yet Tr-d10 showed faint expression and did not co-localize with NF200, suggesting limited interaction between the tracer and spinal nerve pathways (**Suppl. 3c**). Similarly, the distribution of another tracer, OVA488, was absent in peripheral nerves but present in the spinal cord and dorsal root ganglion in prox1-Tom mice (Suppl. 3d). No significant prox1-Tom positive lymphatic vessels were observed along the spinal nerve. The lack of lymphatic vessels may not allow lymphatic absorption of CSF at this site. Instead, as a positive control (10), the optic nerve is encircled by OVA488 with prox1-Tom delineating the optic nerve as the optic nerve expresses prox1 (52) (**Suppl. 3d**).

Moreover, Tr-d10 was not detected in SLN (**Suppl. 4**), indicating that SLN may not be a significant CSF drainage pathway following ICM injection. The absence of Tr-d10 in SLN after ICM injection, compared to its presence following footpad injection of the tracer (**Suppl. 4**), raises questions about the role of sacral lymph nodes in CSF drainage. These results for the two alternative CSF drainage routes may be attributed to the compensatory function of the blood vessels’ drainage of CSF in the nasal mucosa for the disrupted lymphatic function.

In mice, the CLNs comprise sCLNs and dCLNs. The precise connectivity between these two lymph node groups remains undefined. Previous mouse studies focusing on lymphatic CSF drainage disruption typically involved ligation of only the dCLNs (28, 53). This situation assumes that the efferent lymphatic vessels from the sCLNs feed into the dCLNs. Contrary to this assumption, our findings reveal that the efferent vessels of the sCLNs and the dCLNs run in parallel without any direct linkage between the two sets of lymph nodes, especially concerning their efferent pathways. Hence, to effectively interrupt the CSF drainage function of the cervical lymphatic system, it may be crucial to simultaneously ligate both lymph node groups, as one may compensate for the disruption of the other. In our experiment, we achieved complete obstruction of the CSF’s cervical lymphatic drainage by ligating both lymph node groups, targeting a critical pathway for CSF drainage (2, 12, 17). Despite ligating lymph nodes, we noted CSF tracer clearance from the brain and consistent tracer levels in the urine, highlighting the nasal mucosal blood pathway as an effective alternative for CSF drainage. This finding underscores that while lymphatic CSF clearance is vital, it is not solely crucial for CSF management. In our extended study, young wild-type mice (aged 2 months) managed to survive beyond 20 months post bilateral ligation of both sCLNs and dCLNs, showing no significant variance from the control group (data not shown). Moreover, there is no documented rise in AD incidence post-CLN removal in humans, which could be indicative of cancer treatments, suggesting that the critical role attributed to CLNs may be overstated.

Furthermore, an analysis of the absorption capabilities of lymphatic vessels versus blood capillaries for ISF reveals distinctions that are not merely quantitative in terms of capacity but also qualitative in nature (4, 10, 22, 28). It is widely accepted that small soluble molecules predominantly get absorbed by blood capillaries, whereas larger protein molecules are more apt to be taken up by lymphatic vessels, a distinction that stems from the differing permeabilities of these two capillary types (blood and lymph) (22, 28). Blood capillaries possess a more ’closed’ architecture, restricting the entry of larger molecules in contrast to the lymphatic capillaries. The latter exhibit significantly higher permeability due to their incomplete basolateral membranes and the absence of inter-endothelial tight junctions (54), facilitating the free passage of macromolecules from the interstitial space. However, the data regarding the degree of macromolecule permeability through the capillary blood vessel wall are ambiguous.

Studies have demonstrated that macromolecules with a MW spectrum ranging from 5.6 to 66 kDa can traverse the endothelium of blood capillaries via the transcellular pathway (55, 56). In our study, we employed the Tr-d10 tracer, with a molecular weight (MW) of 10,000 (57), which is close to and slightly larger than the MW of Aβ (45,00). Aβ is considered to be the culprit in the development of AD, and the failure of Aβ clearance is believed to be an important factor leading to AD (2, 4, 28). Thus, Tr-d10 serves as a proxy for Aβ removal dynamics in CSF; the tracer’s distinct visibility under bright field illumination aids in tracing CSF drainage routes. Additionally, we utilized a higher MW tracer, OVA555 (MW 45,000, based on ovalbumin), and noted no discernible difference in the clearance rates of both tracers, even with disrupted cervical lymphatic drainage. Furthermore, our observations revealed that the high-MW tracer was also directed to the facial vein (**Suppl. 5**). These findings indicate the high permeability of the vasculature beneath the nasal mucosa, capable of permitting the passage of even large protein molecules. This phenomenon partially clarifies why impairing the CSF drainage through cervical lymphatics does not intensify the buildup of Aβ or exacerbate Aβ-associated pathological alterations in AD mice.

To validate our hypothesis, we examined if the drainage dynamics from the mice’s hind legs mirrored that of CSF drainage. Our study delved into the efficiency of tracer drainage following injections into the mouse footpad, a conventional technique for evaluating the lymphatic system’s effectiveness. This approach has been regularly employed to gain insights into the lymphatic system’s role in drainage management (58–60) (**Suppl. 6a and 6b**). However, the potential for substances injected into the footpad to be cleared via blood vessels remained unexamined. Our research verified that tracers injected into the footpad could effectively be transported through the lymphatic system to the PLN and ILN (**Suppl. 6c and 6d**), aligning with previous observations (60). Interestingly, we also discovered that the tracer visibly filled the femoral veins on the injected side but was absent on the contralateral side at all observed timepoints (**Suppl. 7**). This observation indicates a direct blood vessel drainage pathway for the tracer. Moreover, even after ligating both the PLN and ILN to obstruct the lymphatic route potentially, the tracer continued to be detected in the urine at a rate comparable to that in mice without any lymph node ligation (**Suppl. 8**). This suggests that the blood vessel system can adequately compensate for the impaired lymphatic function, ensuring efficient drainage of substances injected into the footpad. These results are consistent with those of Kagan et al. (56). Their study utilized three types of macromolecules, including bovine insulin, bovine serum albumin, and recombinant human erythropoietin alpha, with molecular weights from 5.6 to 66 kDa. Upon subcutaneous administration, it was discovered that the absorption of all tested macromolecules predominantly occurred through direct penetration into blood capillaries, with the lymphatic system contributing minimally (approximately 3%) to the absorption process, regardless of the broad spectrum of MW involved. Although the results of others (61, 62) differ from this, these differences only reflect variations in the proportion of subcutaneous drug absorption by lymphatic vessels. However, all these results underscore the undeniable fact of blood capillaries’ absorption of subcutaneous drugs. These results again support the possibility of blood vessels’ absorption of CSF in nasal mucosa.

Our research has elucidated previously unchartered territories in understanding the CSF’s circulatory dynamics, broadening the scope of the established theories about the lymphatic system and its relation to the CNS. The most striking discovery from the present study has been identifying blood vessels in the nasal mucosa involved in CSF drainage. This finding holds profound implications for how we perceive the CNS’s drainage mechanism and its interplay with the lymphatic system. Taken together, the results from our study compel us to reconsider the vascular organization around the nasal mucosa. Understanding these pathways is not just an academic exercise; it has tangible implications for medical science. If the blood vessels in the nasal mucosa serve as a conduit for CSF drainage, it could herald innovative approaches to treating brain disorders. In conclusion, the shifting understanding from lymphatic to blood vessel pathways, combined with the complex dynamics of blood vessels in CSF drainage, underscores the necessity for revisiting and updating our knowledge about this critical anatomical region. As with any transformative study, our findings also open doors for further investigations to corroborate these observations and delve deeper into these revised vascular pathways’ physiological and pathological implications.

## Acknowledgements

This work was supported by research funds from the Health@InnoHK Program launched by the Innovation Technology Commission of the Hong Kong SAR, China.

## Author contributions

Yuan Q performed the experiments, analyzed data, and wrote the paper; Satyanarayanan SK, Lee MY, Wang Y, Xian XF, He L, Song Y, and Wu W revised and wrote the manuscript; Yan L and Zhou Y performed the animal experiments; and Su H, Lin ZX and Qin D supervised the entire project, critically involved in study design, interpretation of the data and manuscript revision. All authors contributed to and approved the final manuscript.

## Supplementary Methods

### a) Tracer drainage in the spinal cord and optic nerve

The spinal cord extending from the cervical to the lumbar region was exposed. The cervical cord and its spinal nerves were dissected and preserved in a fixative solution containing 4% paraformaldehyde (PFA) before being immersed in a 30% sucrose solution. Subsequently, the spinal cords with their associated spinal nerves were sectioned (10 µm thickness) using a cryostat. To isolate the optic nerve, the ventral part of the skull was removed to expose the optic nerves. The optic nerves were carefully dissected and promptly placed in the same fixative solution for subsequent sectioning, following the above mentioned procedure.

### b) Tracer drainage in popliteal, inguinal, and sacral lymph nodes

The C57BL/6J mice were anesthetized, as mentioned before. Then, 5 µL of tracers (EB or Tr-d10) were injected into their left footpads. After 30 and 120 mins of the post-injection, mice were euthanized, as mentioned before, and dissected to locate the LNs of interest [popliteal (PLN), inguinal (ILN), and sacral lymph nodes (SLN)]. Similarly, the nodes were extracted and processed as previously mentioned.

---

**Suppl. 1:**
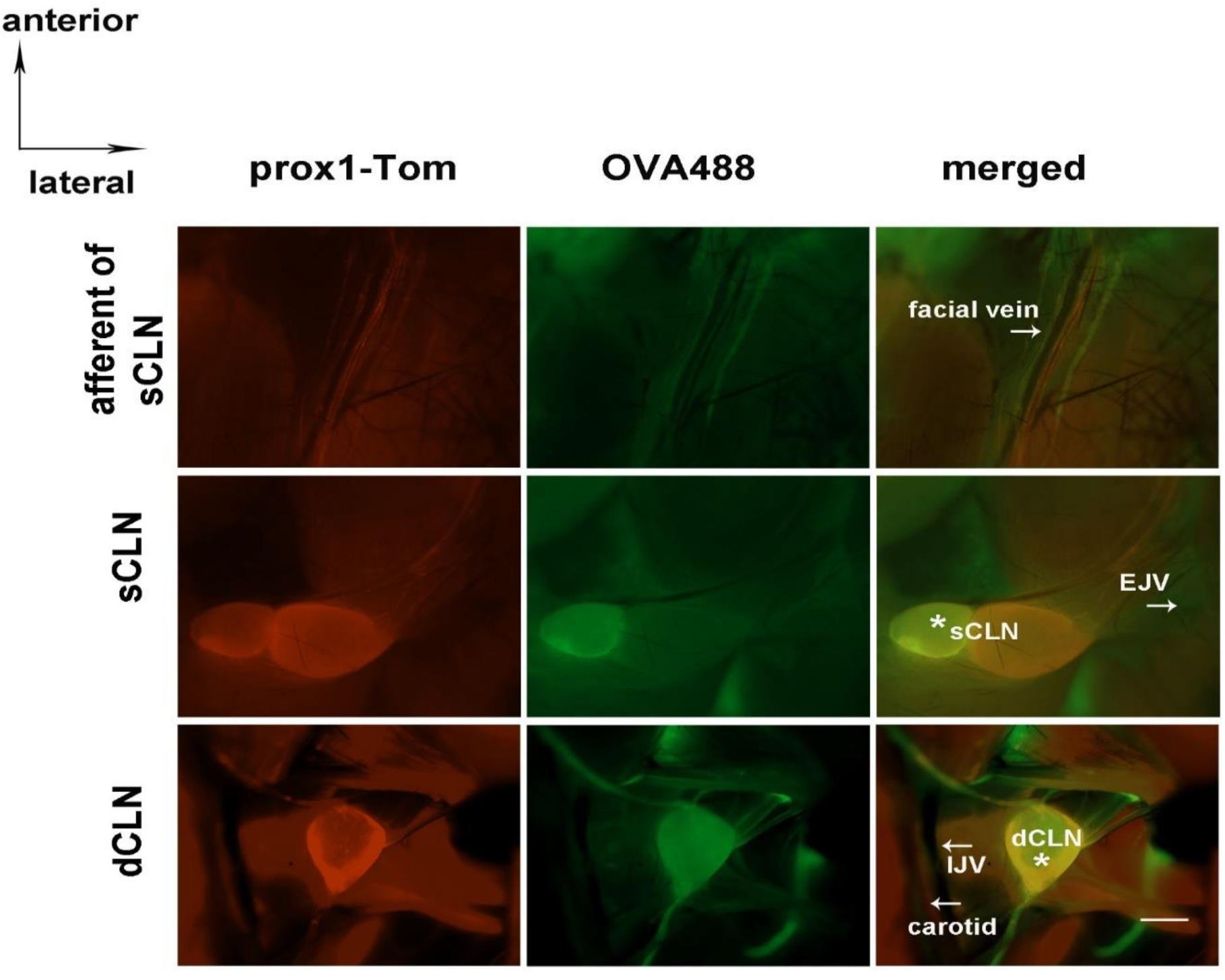
In Vivo Visualization of CSF Tracer in Prox1 tdTom Mice. Representative images showing sCLNs and dCLN with their afferent and efferent vessels after ICM injection of OVA488 in Prox1 tdTom mice. The tracer OVA488 is observed within the cervical lymphatic structures of the mice (stars). No tracer is observed in the facial veins and jugular veins. Scale bar represents 1 mm.

**Suppl. 2:**
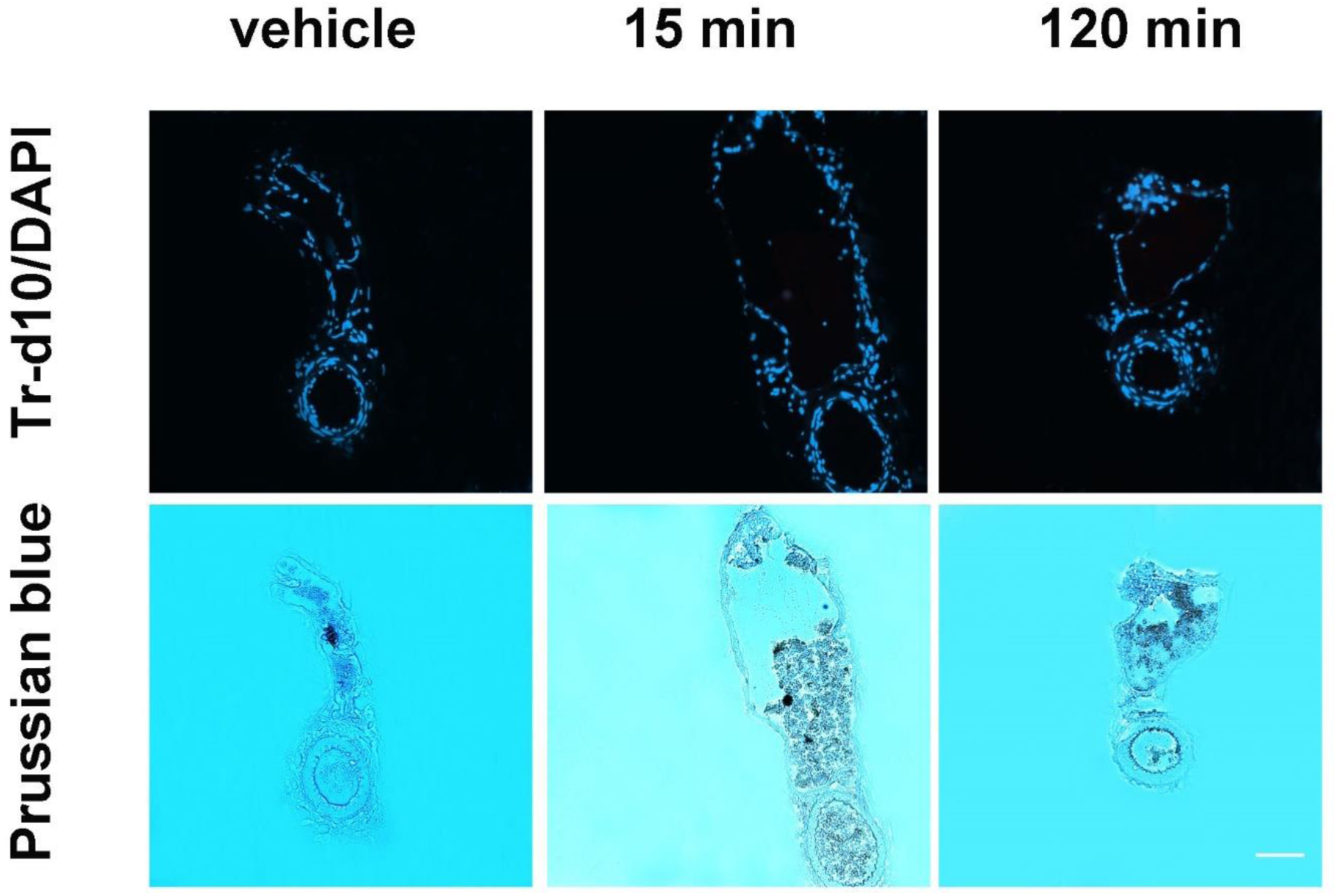
CSF Tracer Distribution in femoral arteries and veins. Representative images showing femoral vein and artery after ICM injection of Tr-d10 or vehicle in the WT mice. CSF tracer Tr-d10 is not observed in the femoral vein and artery of the mice at different time points (15 and 120 min) as in the vehicle group. Cell nuclei are counterstained with DAPI. The corresponding Prussian blue staining of the same sections at the respective time points identifies iron within blood vessels, providing a morphological context to the fluorescence images. Scale bar represents 40 µm.

**Suppl. 3:**
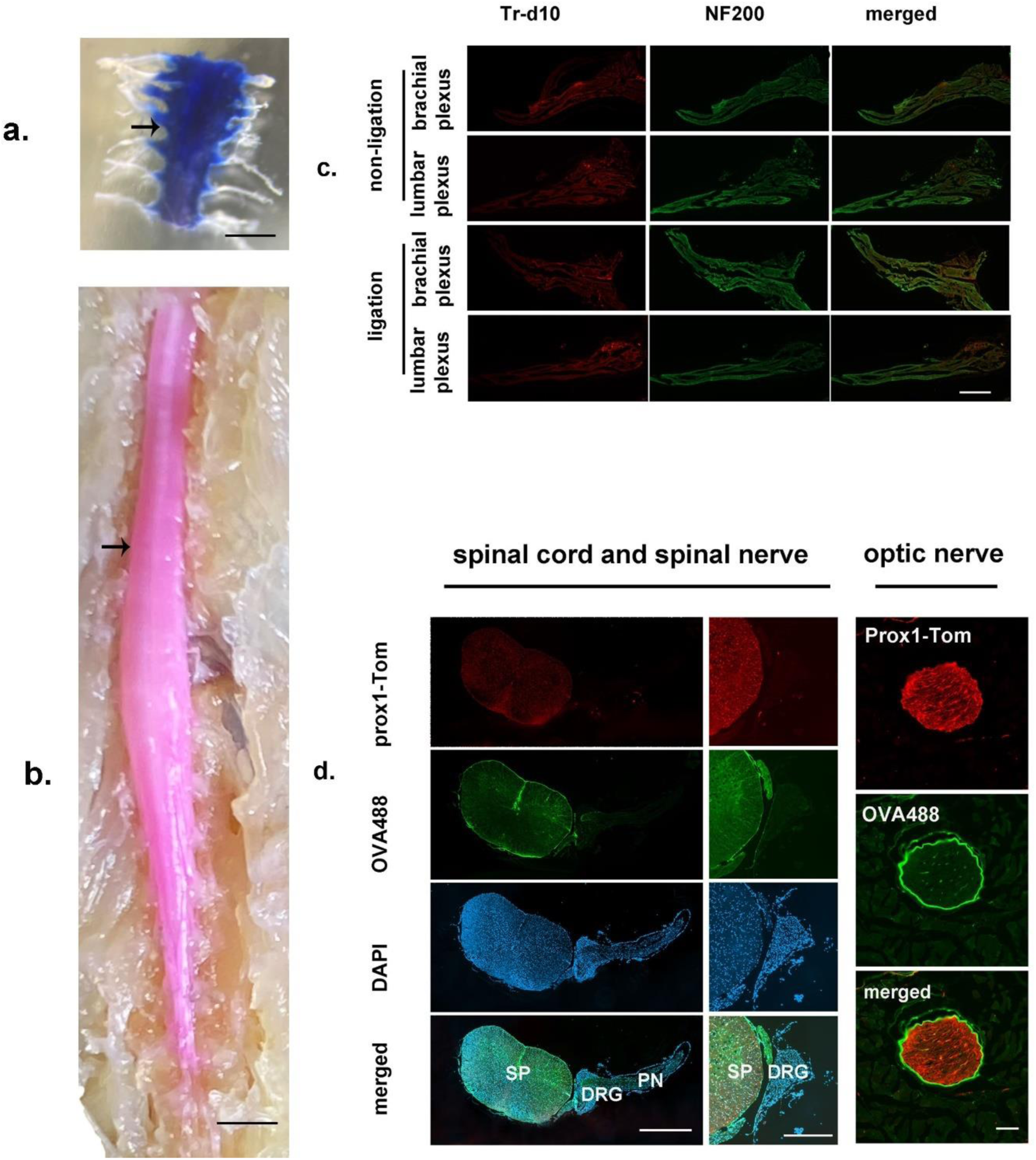
CSF Tracer Distribution in the Spinal Nerve Pathway. (a) & (b) CSF tracer EB and Tr-d10 are only observed within the spinal subarachnoid space (arrows) but not in the spinal nerves following ICM injection of either EB or Tr0d10. (c) Representative immunofluorescence images depicting brachial and lumbar plexus in non-ligated and ligated models, with neurofilament 200 (NF200) highlighting nerve fibers following ICM injection of Trd10. No significant Tr-d10 is observed in the spinal nerves, regardless of ligation. (d) Distinct compartmentalization of OVA488 in the spinal cord and dorsal root ganglion (DRG) but not peripheral nerves (PN), as shown in prox1-Tom mice following ICM injection of OVA488. In contrast, as a positive control (*1*) the optic nerve is encircled by OVA488 with prox1-Tom delineating the optic nerve as optic nerve expresses prox1 (*2*). Cell nuclei are counterstained with DAPI. The scale bar in (a) and (b) represents 1 mm. Scale bar in (c) represents 40 µm. Scale bar in the first collum of (d) represents 500 µm. Scale bar in the second collum of (d) represents 200 µm. Scale bar in the third collum of (d) represents 20 µm.

**Suppl. 4:**
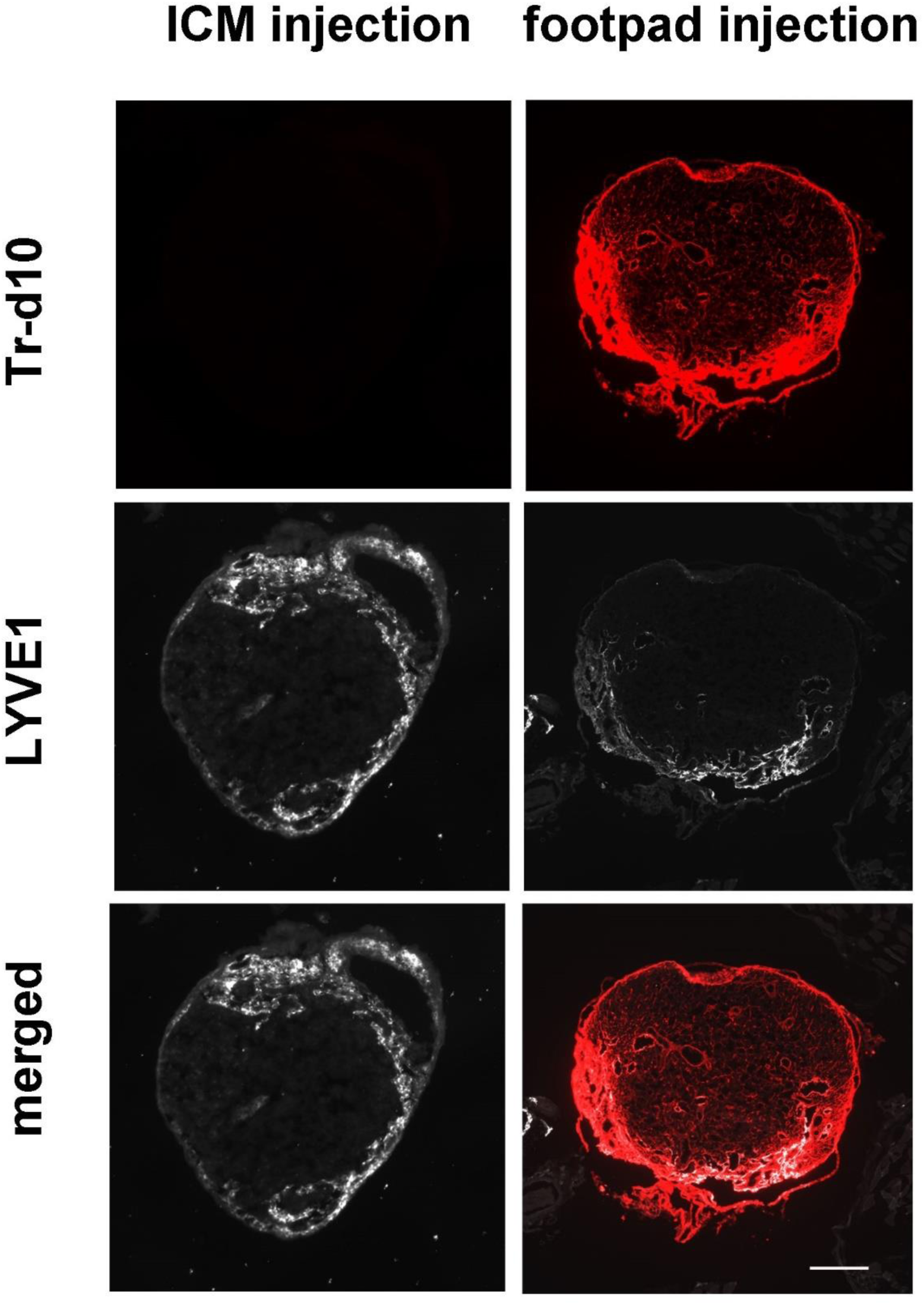
The CSF Tracer Absence in Sacral Lymphatic Nodes (SLN) Post-ICM Injection. A comparison of the tracer Tr-d10 distribution in popliteal lymph nodes (PLN) following ICM and footpad injection of Tr-d10. The footpad injection of Tr-d10 results in the presence of the tracer in the PLN. In contrast, ICM injection of the tracer shows no detectable presence of Tr-d10 in the PLN, indicating that the CSF tracer may not be drained via the PLN. The presence of lymphatic vessel endothelial hyaluronan receptor 1 (LYVE-1) seen in the SLN and PLN highlights the structure of the lymphatic nodes. Scale bar represents 250 µm.

**Suppl. 5:**
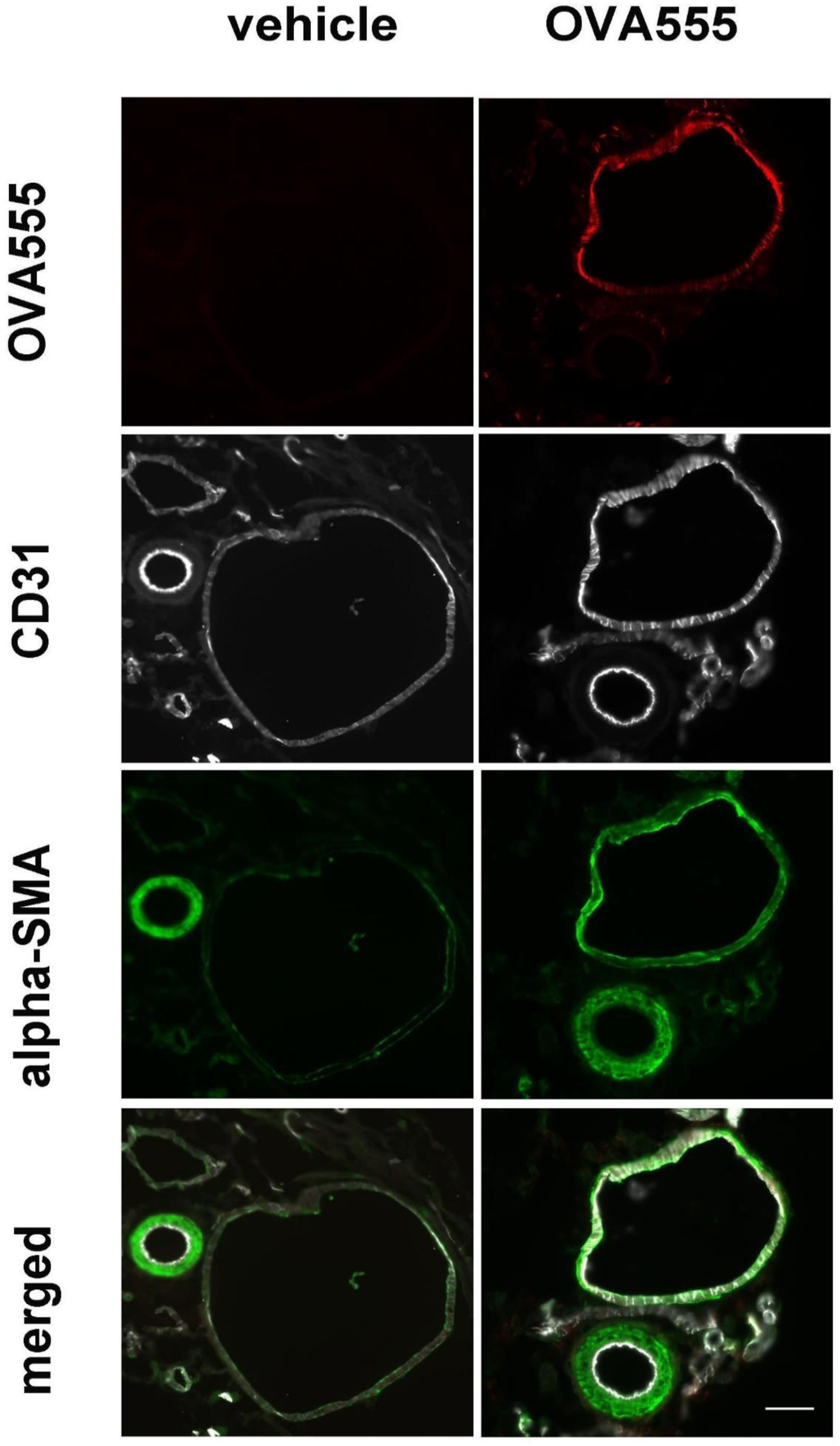
The Drainage of a CSF Tracer with a Larger Molecular Weight. Representative immunofluorescence images depicting the distribution of a CSF tracer with a higher molecular weight (OVA555, 45 kDa) in the facial vein from exsanguinated mice after ICM injection compared to the vehicle. Post-ICM injection of OVA555 results in the presence of the tracer in the facial vein but not the facial artery as evidenced by OVA555+/CD31+/αSMA + immunostaining. The scale bar represents 40 µm.

**Suppl. 6:**
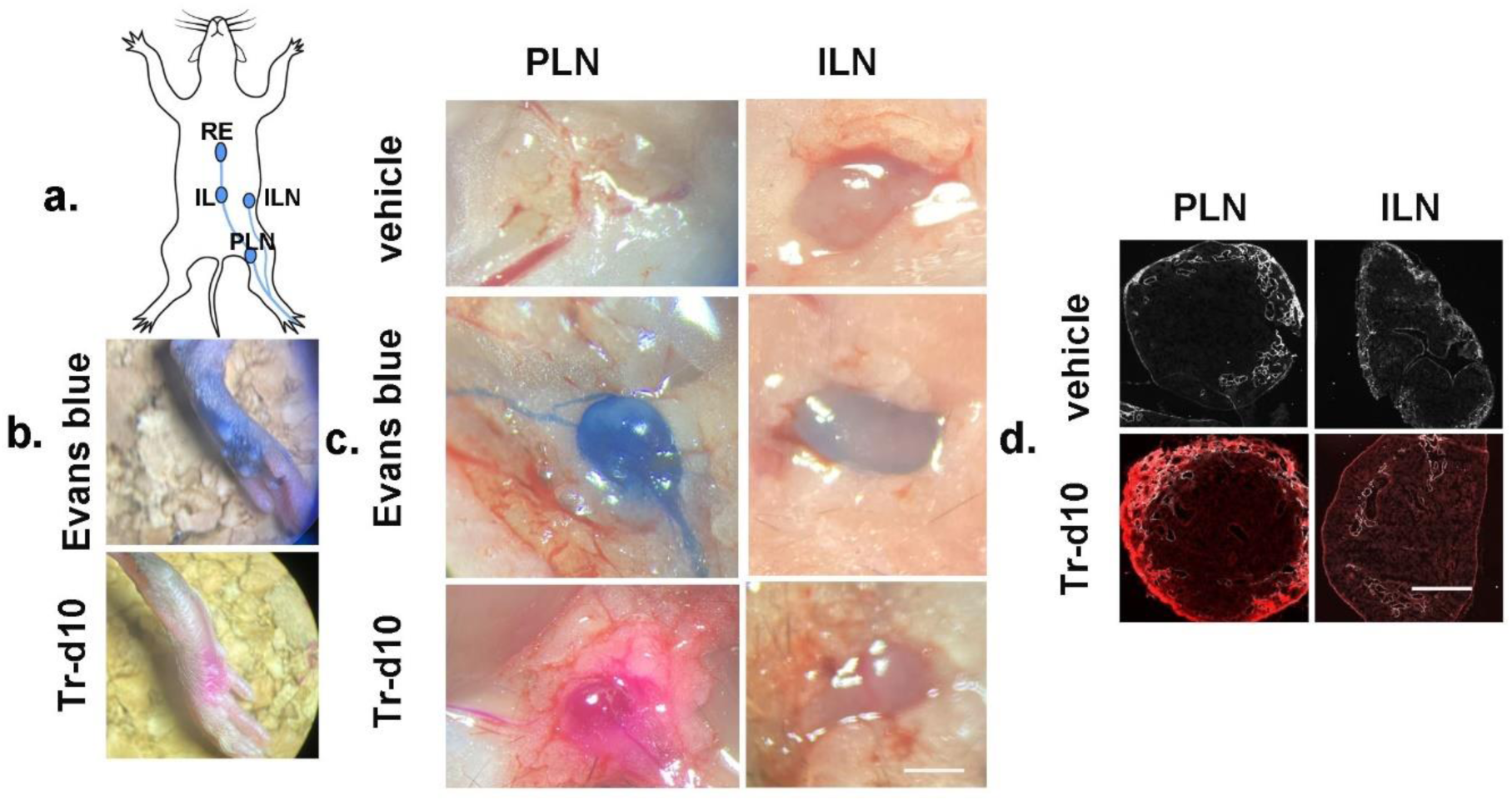
Lymphatic Drainage of the Tracer Tr-d10 Following Footpad Injection of Tr-d10. (a) Diagram illustrating the footpad injection site and the subsequent lymphatic drainage pathways in a mouse model, showing the direction of tracer movement toward the popliteal lymph node (PLN) and inguinal lymph node (ILN). (b) Macroscopic images depicting the visualization of Evans Blue dye (EB) and Tr-d10 after footpad injection. (c) Macroscopic images of PLN and ILN from mice receiving the vehicle, EB, and Tr-d10 injection showing the visible transport of the tracers within the PLN and ILN compared with that in the vehicle group. (d) Representative images depicting the distribution of the tracer Tr-d10 in the PLN and ILN after footpad injection of Tr-d10. The footpad injection of Tr-d10 results in the presence of the tracer in the PLN and ILN. The lymphatic vessel endothelial hyaluronan receptor 1 (LYVE-1) seen in the PLN and ILN highlights the structure of the lymphatic nodes. Scale bar represents 500 µm.

**Suppl. 7:**
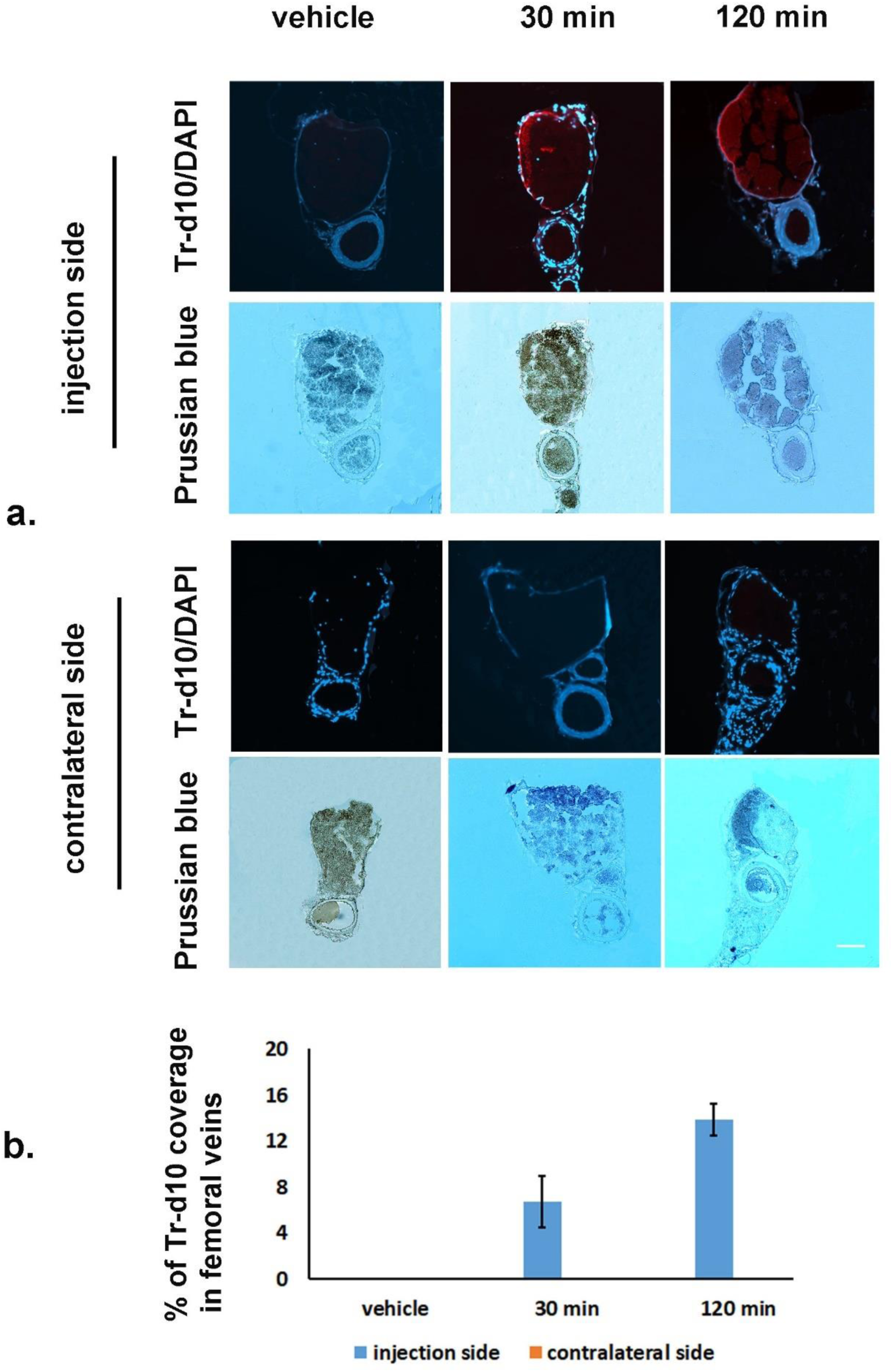
Drainage of the Tracer Tr-d10 via Blood Vessels Following Footpad Injection of Tr-d10. (a) Representative images of the femoral vein and artery in the injection and the contralateral sides following the footpad injection of Tr-d10. A triple staining involving Tr-d10/DAPI/Prussian blue is employed to identify the localization of the tracer Tr-d10. Cell nuclei are counterstained with DAPI, highlighting the femoral vein and artery structure. The presence of the tracer in the veins but not the artery of the injection side at both 30- and 120 minutes post-injection, as well as its absence on the contralateral side, indicates that the tracer in the veins is not indirectly introduced through the systemic circulation but instead directly absorbed via blood capillaries. (b) Quantification of Tr-d10 area fraction within the femoral vein over time (mean ± SEM), demonstrating a peak in tracer presence at 30 minutes and a decrease at 120 minutes. Scale bar represents 40 µm.

**Suppl. 8:**
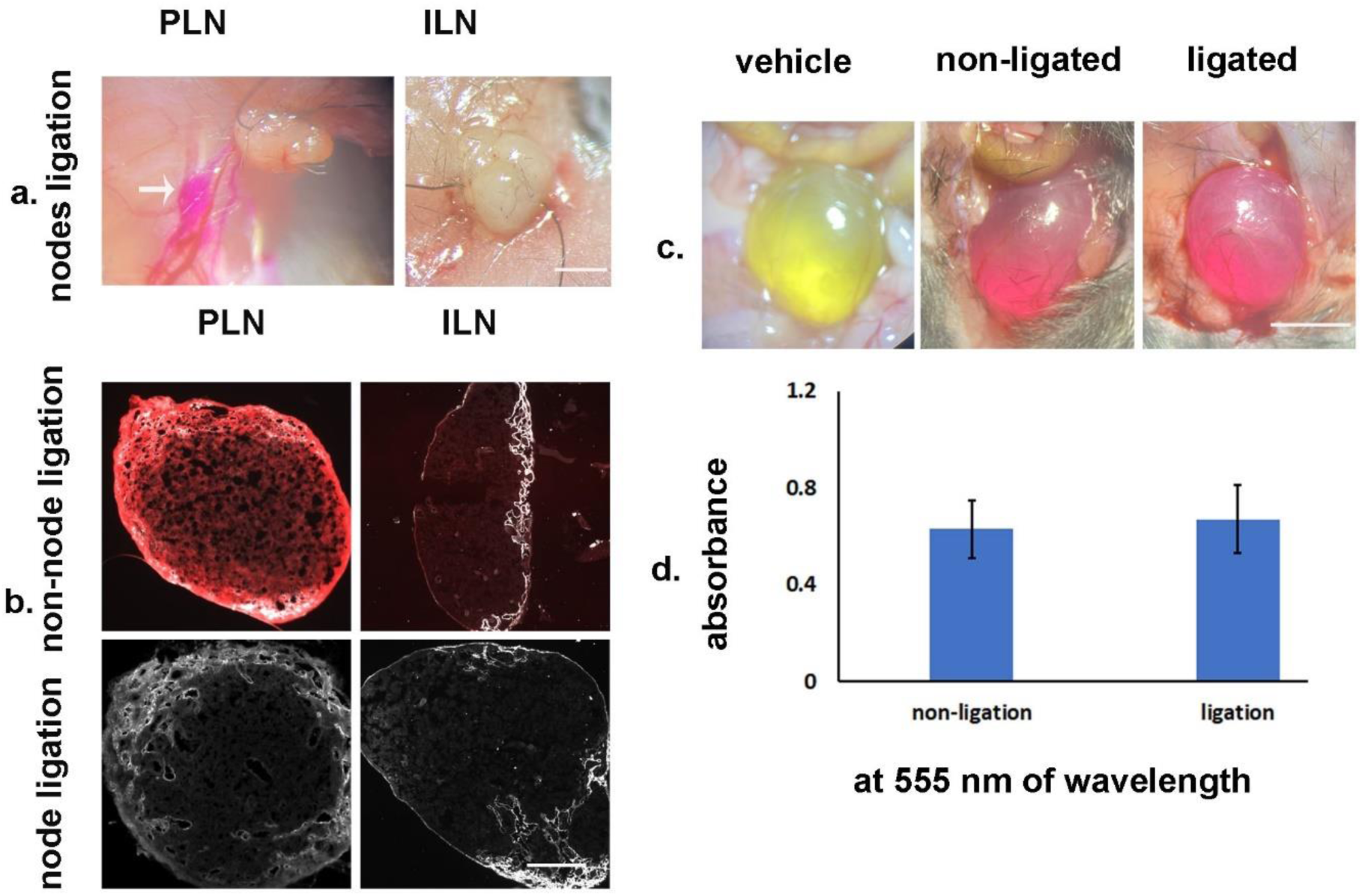
Impact of Popliteal (PLN) and Inguinal (ILN) Lymph Nodes Ligation on Tracer Drainage Following Footpad Injection of the Tr-d10. (a) Macroscopic images of ligated PLN and ILN lymph nodes following footpad injection of Tr-d10. The tracer is visibly present in the afferent lymphatic vessels of the PLN, as shown by an enlargement with accumulated tracer Tr-d10 (an arrow), confirming the interruption of lymphatic flow. (b) Representative lymph nodes’ sections stained with LYVE1 immunostaining after Tr-d10 footpad injection. The non-ligated displayed the presence of Tr-d10 in PLN and ILN following the footpad injection, whereas ligated nodes showed no Tr-d10 signal, indicating that ligation prevented the entry of the tracer. The presence of LYVE-1 seen in the PLN and ILN highlights the structure of the lymphatic nodes. (c) A representative dissected lower abdomen showing urinary bladder at 2 h following Tr-d10 or vehicle footpad injection, with non-ligation and ligation of both PLN and ILN ligation. The presence of pink coloration indicates the tracer’s passage into the urine. (d) Quantitative analysis of Tr-d10 in urine samples based on absorbance at a wavelength of 555 nm. The graph depicts similar absorbance levels for non-ligated and ligated groups, suggesting unaffected tracer elimination from the footpad into the urine following both PLN and ILN ligation. The scale bar in (a) represents 1 mm. Scale bar in (b) represents 400 µm. Scale bar in (c) represents 4 mm.

## References

1. H. H. Damkier, P. D. Brown, J. Praetorius, Cerebrospinal fluid secretion by the choroid plexus. Physiol Rev 93, 1847–1892 (2013).

2. M. Nedergaard, Neuroscience. Garbage truck of the brain. Science 340, 1529–1530 (2013).

3. N. A. Jessen, A. S. Munk, I. Lundgaard, M. Nedergaard, The Glymphatic System: A Beginner’s Guide. Neurochem Res 40, 2583–2599 (2015).

4. S. T. Proulx, Cerebrospinal fluid outflow: a review of the historical and contemporary evidence for arachnoid villi, perineural routes, and dural lymphatics. Cell Mol Life Sci 78, 2429–2457 (2021).

5. I. Spera, S. T. Proulx, A nasal hub for cerebrospinal fluid clearance. Nature Cardiovascular Research 3, 98–99 (2024).

6. Y. Zhou et al., Impaired peri-olfactory cerebrospinal fluid clearance is associated with ageing, cognitive decline and dyssomnia. eBioMedicine 86, (2022).

7. S. Da Mesquita, Z. Fu, J. Kipnis, The Meningeal Lymphatic System: A New Player in Neurophysiology. Neuron 100, 375–388 (2018).

8. S. Da Mesquita et al., Functional aspects of meningeal lymphatics in ageing and Alzheimer’s disease. Nature 560, 185–191 (2018).

9. A. Louveau et al., CNS lymphatic drainage and neuroinflammation are regulated by meningeal lymphatic vasculature. Nat Neurosci 21, 1380–1391 (2018).

10. Q. Ma, B. V. Ineichen, M. Detmar, S. T. Proulx, Outflow of cerebrospinal fluid is predominantly through lymphatic vessels and is reduced in aged mice. Nat Commun 8, 1434 (2017).

11. D. Raper, A. Louveau, J. Kipnis, How Do Meningeal Lymphatic Vessels Drain the CNS? Trends Neurosci 39, 581–586 (2016).

12. I. Spera et al., Open pathways for cerebrospinal fluid outflow at the cribriform plate along the olfactory nerves. EBioMedicine 91, 104558 (2023).

13. S. Kida, A. Pantazis, R. O. Weller, CSF drains directly from the subarachnoid space into nasal lymphatics in the rat. Anatomy, histology and immunological significance. Neuropathology and applied neurobiology 19, 480–488 (1993).

14. A. Zakharov, C. Papaiconomou, J. Djenic, R. Midha, M. Johnston, Lymphatic cerebrospinal fluid absorption pathways in neonatal sheep revealed by subarachnoid injection of Microfil. Neuropathology and applied neurobiology 29, 563–573 (2003).

15. M. Johnston, A. Zakharov, L. Koh, D. Armstrong, Subarachnoid injection of Microfil reveals connections between cerebrospinal fluid and nasal lymphatics in the non-human primate. Neuropathology and applied neurobiology 31, 632–640 (2005).

16. Y. Decker et al., Magnetic resonance imaging of cerebrospinal fluid outflow after low-rate lateral ventricle infusion in mice. JCI Insight 7, (2022).

17. L. Jacob et al., Conserved meningeal lymphatic drainage circuits in mice and humans. J Exp Med 219, (2022).

18. J. H. Ahn et al., Meningeal lymphatic vessels at the skull base drain cerebrospinal fluid. Nature 572, 62–66 (2019).

19. J. H. Yoon et al., Nasopharyngeal lymphatic plexus is a hub for cerebrospinal fluid drainage. Nature 625, 768–777 (2024).

20. H. Wiig, M. A. Swartz, Interstitial fluid and lymph formation and transport: physiological regulation and roles in inflammation and cancer. Physiol Rev 92, 1005–1060 (2012).

21. Y. Himeno, M. Ikebuchi, A. Maeda, A. Noma, A. Amano, Mechanisms underlying the volume regulation of interstitial fluid by capillaries: a simulation study. Integr Med Res 5, 11–21 (2016).

22. A. G. Warren, H. Brorson, L. J. Borud, S. A. Slavin, Lymphedema: a comprehensive review. Ann Plast Surg 59, 464–472 (2007).

23. Y. Watanabe et al., Development and Themes of Diagnostic and Treatment Procedures for Secondary Leg Lymphedema in Patients with Gynecologic Cancers. Healthcare (Basel) 7, (2019).

24. V. Angeli, H. Y. Lim, Biomechanical control of lymphatic vessel physiology and functions. Cellular & Molecular Immunology 20, 1051–1062 (2023).

25. H. F. Cserr, C. J. Harling-Berg, P. M. Knopf, Drainage of brain extracellular fluid into blood and deep cervical lymph and its immunological significance. *Brain pathology (Zurich*, Switzerland) 2, 269–276 (1992).

26. A. Aspelund et al., A dural lymphatic vascular system that drains brain interstitial fluid and macromolecules. J Exp Med 212, 991–999 (2015).

27. A. Louveau et al., Structural and functional features of central nervous system lymphatic vessels. Nature 523, 337–341 (2015).

28. L. Wang et al., Deep cervical lymph node ligation aggravates AD-like pathology of APP/PS1 mice. *Brain pathology (Zurich*, Switzerland) 29, 176–192 (2019).

29. M. E. Smith, D. G. Morton, in The Digestive System (Second Edition), M. E. Smith, D. G. Morton, Eds. (Churchill Livingstone, 2010), pp. 107–127.

30. D. G. Potts, V. Deonarine, W. Welton, Perfusion studies of the cerebrospinal fluid absorptive pathways in the dog. Radiology 104, 321–325 (1972).

31. A. Zakharov, C. Papaiconomou, M. Johnston, Lymphatic vessels gain access to cerebrospinal fluid through unique association with olfactory nerves. Lymphat Res Biol 2, 139–146 (2004).

32. R. Mollanji, R. Bozanovic-Sosic, A. Zakharov, L. Makarian, M. G. Johnston, Blocking cerebrospinal fluid absorption through the cribriform plate increases resting intracranial pressure. Am J Physiol Regul Integr Comp Physiol 282, R1593–1599 (2002).

33. S. Yamada et al., MRI tracer study of the cerebrospinal fluid drainage pathway in normal and hydrocephalic guinea pig brain. Tokai J Exp Clin Med 30, 21–29 (2005).

34. B. A. Walter, V. A. Valera, S. Takahashi, T. Ushiki, The olfactory route for cerebrospinal fluid drainage into the peripheral lymphatic system. Neuropathology and applied neurobiology 32, 388–396 (2006).

35. M. Johnston, A. Zakharov, C. Papaiconomou, G. Salmasi, D. Armstrong, Evidence of connections between cerebrospinal fluid and nasal lymphatic vessels in humans, non-human primates and other mammalian species. Cerebrospinal Fluid Res 1, 2 (2004).

36. G. Nagra, L. Koh, A. Zakharov, D. Armstrong, M. Johnston, Quantification of cerebrospinal fluid transport across the cribriform plate into lymphatics in rats. Am J Physiol Regul Integr Comp Physiol 291, R1383–1389 (2006).

37. L. A. Murtha et al., Cerebrospinal fluid is drained primarily via the spinal canal and olfactory route in young and aged spontaneously hypertensive rats. Fluids Barriers CNS 11, 12 (2014).

38. B. Bedussi et al., Clearance from the mouse brain by convection of interstitial fluid towards the ventricular system. Fluids Barriers CNS 12, 23 (2015).

39. S. Kwon, C. F. Janssen, F. C. Velasquez, E. M. Sevick-Muraca, Fluorescence imaging of lymphatic outflow of cerebrospinal fluid in mice. J Immunol Methods 449, 37–43 (2017).

40. J. N. Norwood et al., Anatomical basis and physiological role of cerebrospinal fluid transport through the murine cribriform plate. Elife 8, (2019).

41. K. T. Householder, S. Dharmaraj, D. I. Sandberg, R. J. Wechsler-Reya, R. W. Sirianni, Fate of nanoparticles in the central nervous system after intrathecal injection in healthy mice. Sci Rep 9, 12587 (2019).

42. M. Brady et al., Cerebrospinal fluid drainage kinetics across the cribriform plate are reduced with aging. Fluids Barriers CNS 17, 71 (2020).

43. P. Papakyriakopoulou, G. Valsami, N. P. E. Kadoglou, Nose-to-Heart Approach: Unveiling an Alternative Route of Acute Treatment. Biomedicines 12, (2024).

44. G. Grevers, [Morphology of the vascular endothelium of the nasal mucosa]. Laryngol Rhinol Otol (Stuttg) 66, 625–628 (1987).

45. G. Grevers, The role of fenestrated vessels for the secretory process in the nasal mucosa: a histological and transmission electron microscopic study in the rabbit. Laryngoscope 103, 1255–1258 (1993).

46. G. Grevers, U. Herrmann, Fenestrated endothelia in vessels of the nasal mucosa. An electron-microscopic study in the rabbit. Arch Otorhinolaryngol 244, 55–60 (1987).

47. L. A. Keller, O. Merkel, A. Popp, Intranasal drug delivery: opportunities and toxicologic challenges during drug development. Drug Deliv Transl Res 12, 735–757 (2022).

48. C. Tucker, L. Tucker, K. Brown, The Intranasal Route as an Alternative Method of Medication Administration. Crit Care Nurse 38, 26–31 (2018).

49. J. B. Pawlak, K. M. Caron, Lymphatic Programing and Specialization in Hybrid Vessels. Front Physiol 11, 114 (2020).

50. M. Mancini et al., Head and Neck Veins of the Mouse. A Magnetic Resonance, Micro Computed Tomography and High Frequency Color Doppler Ultrasound Study. PLoS One 10, e0129912 (2015).

51. S. W. Bothwell, D. Janigro, A. Patabendige, Cerebrospinal fluid dynamics and intracranial pressure elevation in neurological diseases. Fluids Barriers CNS 16, 9 (2019).

52. Y. Yang, L. Zhang, G. Li, L. J. I. O. Chen, V. Science, Novel characterization of Prox-1 expression in the optic nerve. 59, 312–312 (2018).

53. N. Kinota et al., Blockage of CSF Outflow in Rats after Deep Cervical Lymph Node Ligation Observed Using Gd-based MR Imaging. Magn Reson Med Sci, (2023).

54. M. A. Swartz, The physiology of the lymphatic system. Adv Drug Deliv Rev 50, 3–20 (2001).

55. M. Bendayan, Morphological and cytochemical aspects of capillary permeability. Microsc Res Tech 57, 327–349 (2002).

56. L. Kagan et al., The role of the lymphatic system in subcutaneous absorption of macromolecules in the rat model. Eur J Pharm Biopharm 67, 759–765 (2007).

57. J. J. Iliff et al., A paravascular pathway facilitates CSF flow through the brain parenchyma and the clearance of interstitial solutes, including amyloid beta. Sci Transl Med 4, 147ra111 (2012).

58. N. L. Tilney, The systemic distribution of soluble antigen injected into the footpad of the laboratory rat. Immunology 19, 181–184 (1970).

59. Y. Yamaji et al., Development of a mouse model for the visual and quantitative assessment of lymphatic trafficking and function by in vivo imaging. Scientific Reports 8, 5921 (2018).

60. M. I. Harrell, B. M. Iritani, A. Ruddell, Lymph node mapping in the mouse. J Immunol Methods 332, 170–174 (2008).

61. S. A. Charman, A. M. Segrave, G. A. Edwards, C. J. Porter, Systemic availability and lymphatic transport of human growth hormone administered by subcutaneous injection. J Pharm Sci 89, 168–177 (2000).

62. D. N. McLennan, C. J. Porter, S. A. Charman, Subcutaneous drug delivery and the role of the lymphatics. Drug Discov Today Technol 2, 89–96 (2005).

