## Supplementary materials for "Cerebrospinal Fluid Dynamics: Uncovering Alternative Blood Vessel Clearance Mechanisms"

Running title: ***Alternative Blood Vessel Clearance Mechanisms in CSF Dynamics***

*Corresponding authors:

Huanxing Su, State Key Laboratory of Quality Research in Chinese Medicine, Institute of Chinese Medical Sciences, University of Macau, Macao, China


Zhi-Xiu Lin, School of Chinese Medicine, Faculty of Medicine, The Chinese University of Hong Kong, Shatin, New Territory, Hong Kong Special Administration Region, China.


Dajiang Qin, Key Laboratory of Biological Targeting Diagnosis, Therapy and Rehabilitation of Guangdong Higher Education Institutes, The Fifth Affiliated Hospital of Guangzhou Medical University, Guangzhou, China


### Supplementary Methods

#### *a) Tracer drainage in the spinal cord and optic nerve*

The spinal cord extending from the cervical to the lumbar region was exposed. The cervical cord and its spinal nerves were dissected and preserved in a fixative solution containing 4% paraformaldehyde (PFA) before being immersed in a 30% sucrose solution. Subsequently, the spinal cords with their associated spinal nerves were sectioned (10 µm thickness) using a cryostat. To isolate the optic nerve, the ventral part of the skull was removed to expose the optic nerves. The optic nerves were carefully dissected and promptly placed in the same fixative solution for subsequent sectioning, following the above mentioned procedure.

#### *b) Tracer drainage in popliteal, inguinal, and sacral lymph nodes*

The C57BL/6J mice were anesthetized, as mentioned before. Then, 5 µL of tracers (EB or Tr-d10) were injected into their left footpads. After 30 and 120 mins of the post-injection, mice were euthanized, as mentioned before, and dissected to locate the LNs of interest [popliteal (PLN), inguinal (ILN), and sacral lymph nodes (SLN)]. Similarly, the nodes were extracted and processed as previously mentioned.

### Supplementary Figures

**Suppl. 1**

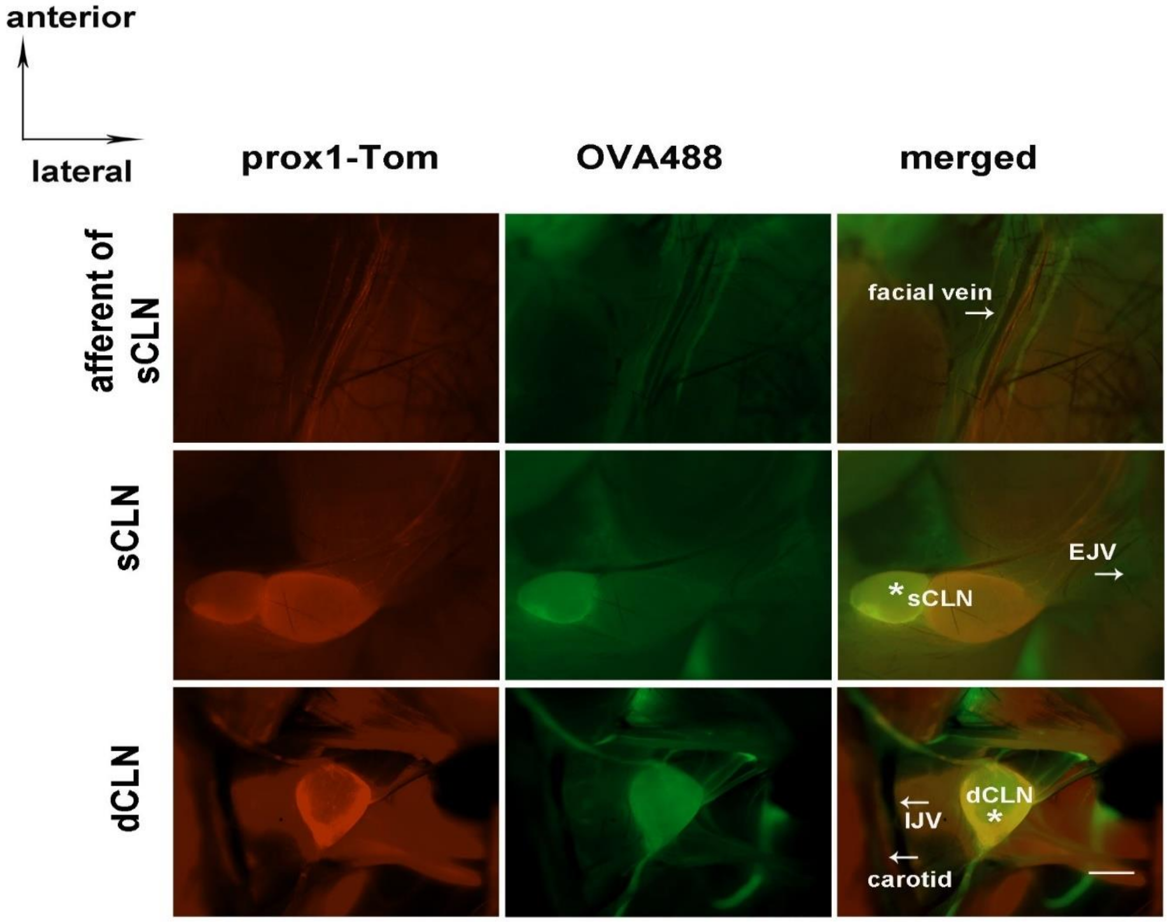

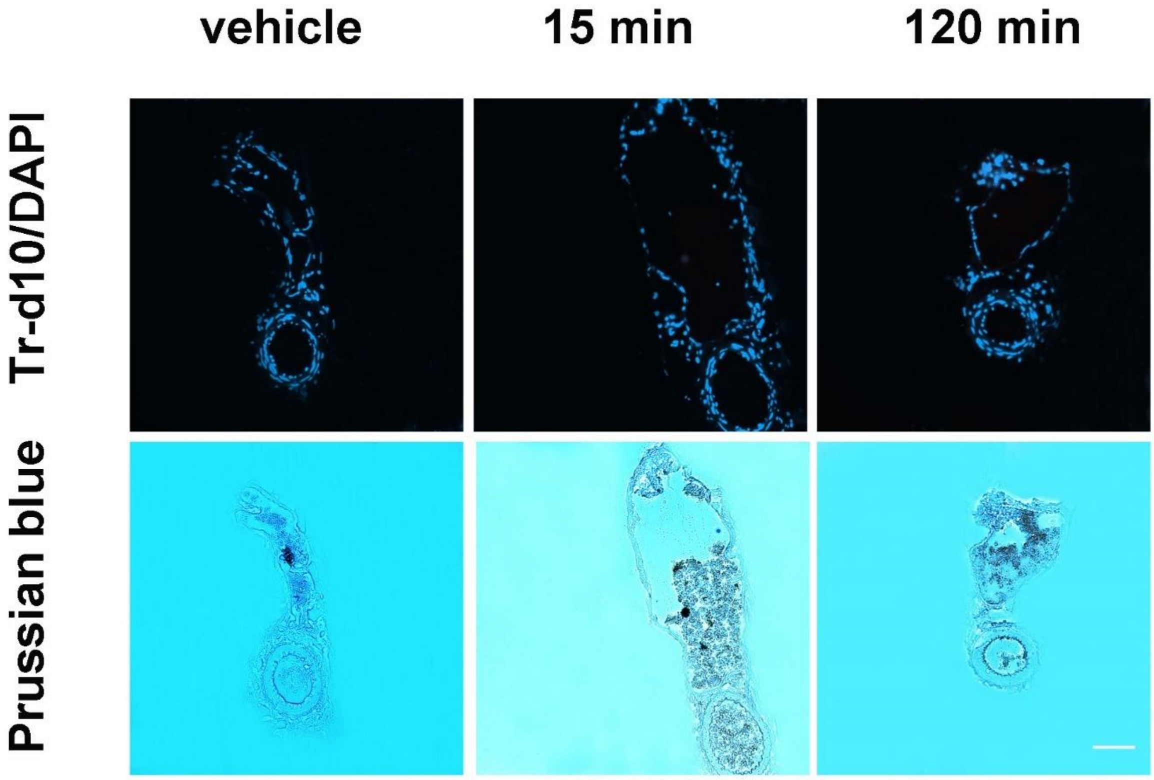
vehicle
15 min
120 min
Tr-d10/DAPI
Prussian blue

Suppl. 3

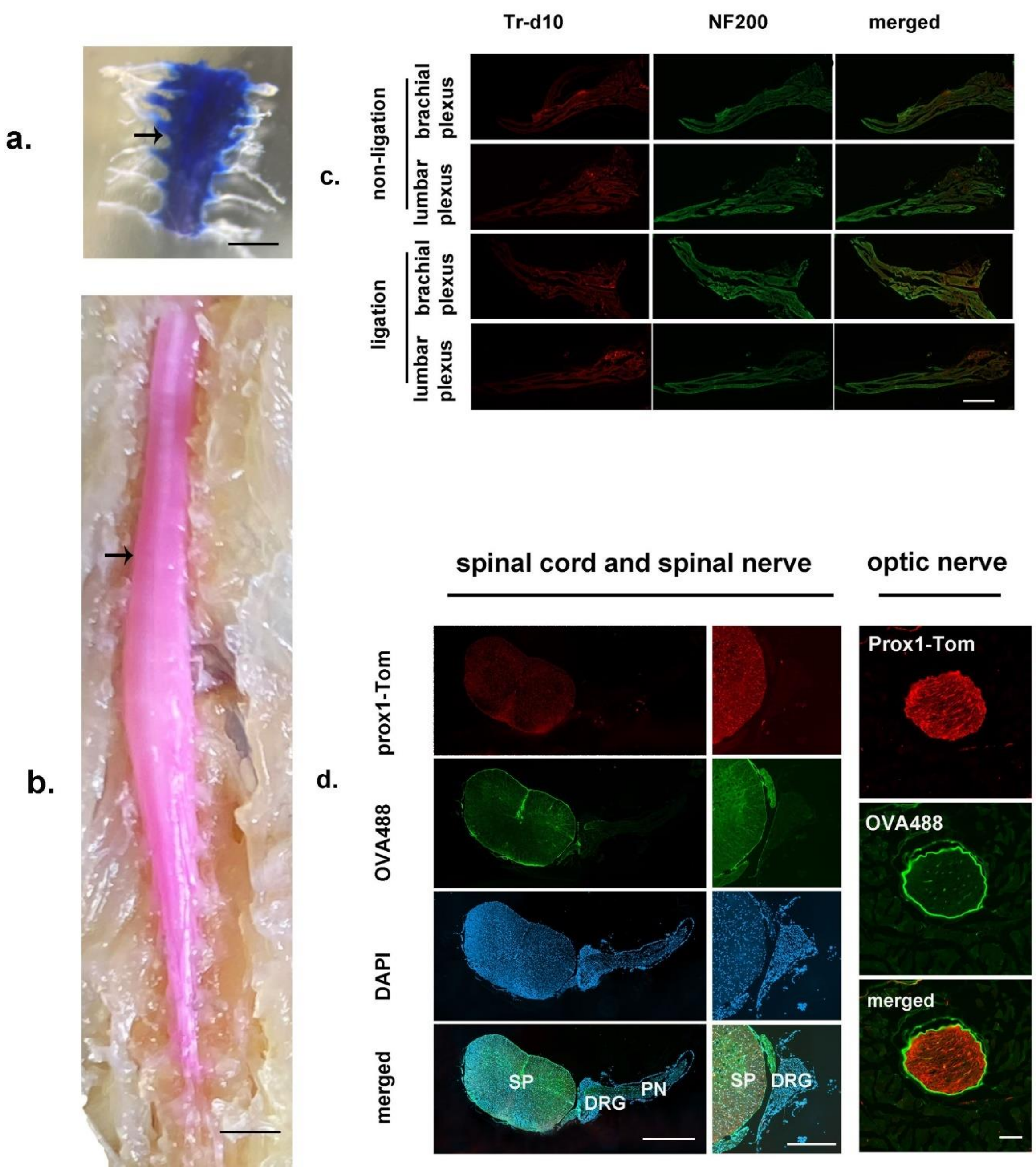
a.
b.
c.
Tr-d10
NF200
merged
non-ligation
brachial plexus
lumbar plexus
ligation
brachial plexus
lumbar plexus
d.
spinal cord and spinal nerve
optic nerve
prox1-Tom
OVA488
DAPI
merged
Prox1-Tom
OVA488
merged
SP
DRG
PN
SP
DRG

**Suppl. 4**

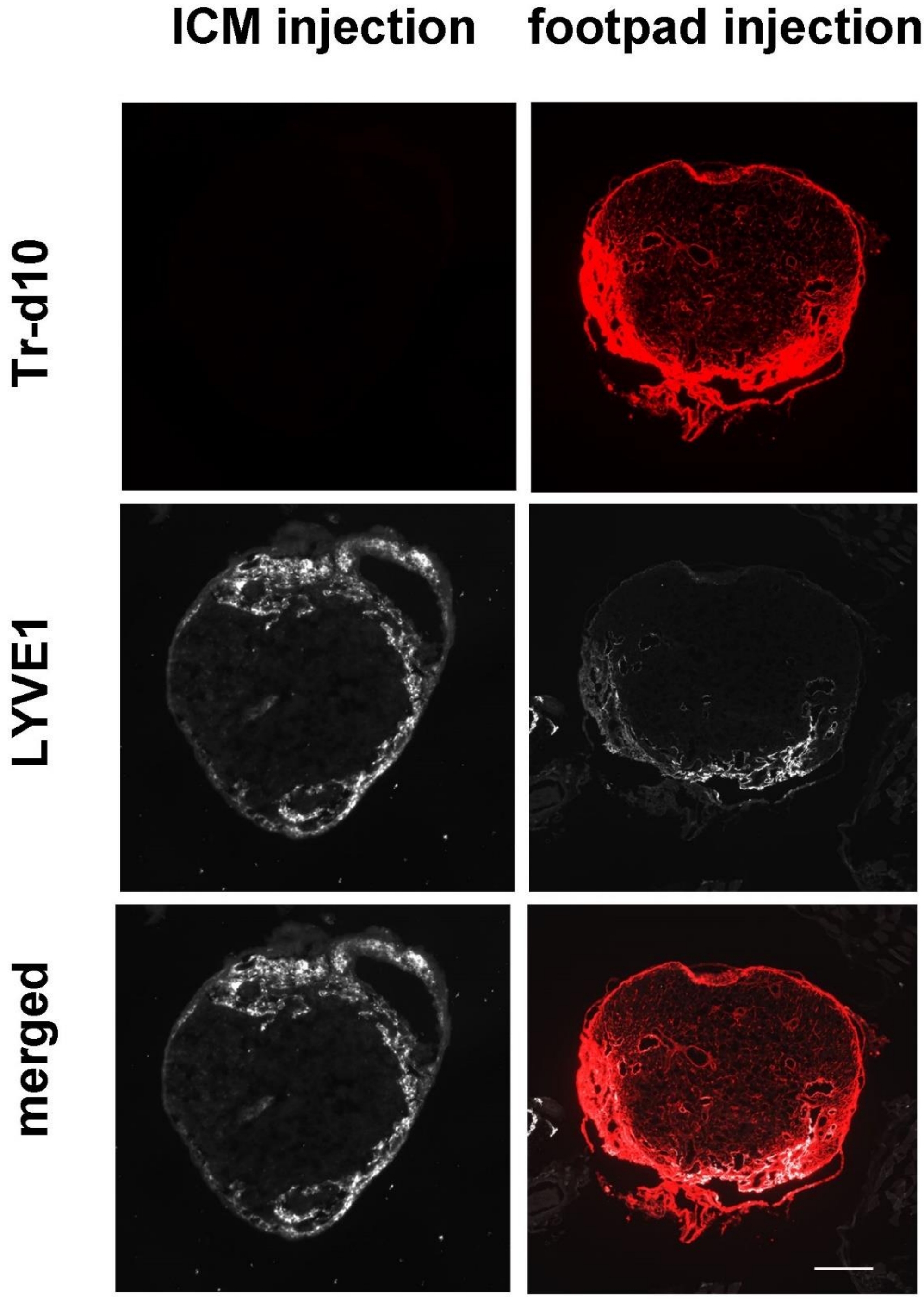

**Suppl. 5**

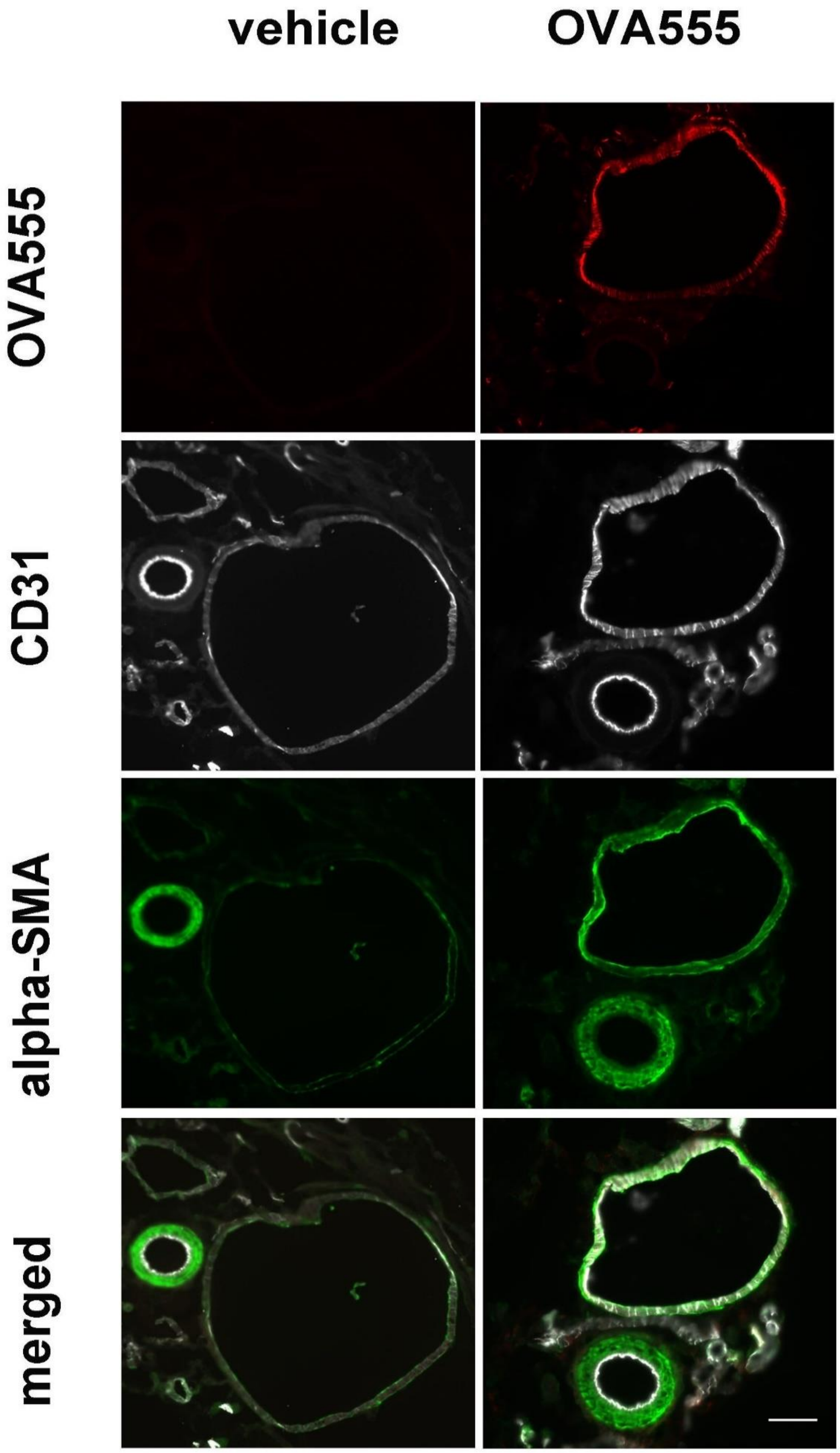

**Suppl. 6**

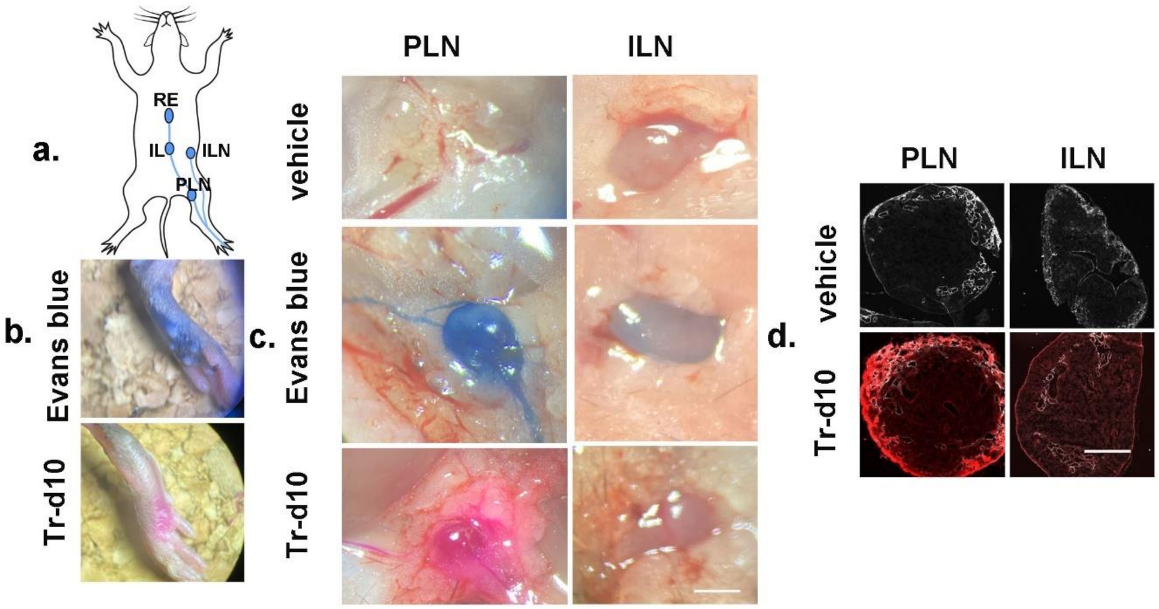

a.
RE
IL
ILN
PLN
b.
Evans blue
Tr-d10
c.
PLN
ILN
vehicle
Evans blue
Tr-d10
d.
PLN
ILN
vehicle
Tr-d10

Suppl. 7

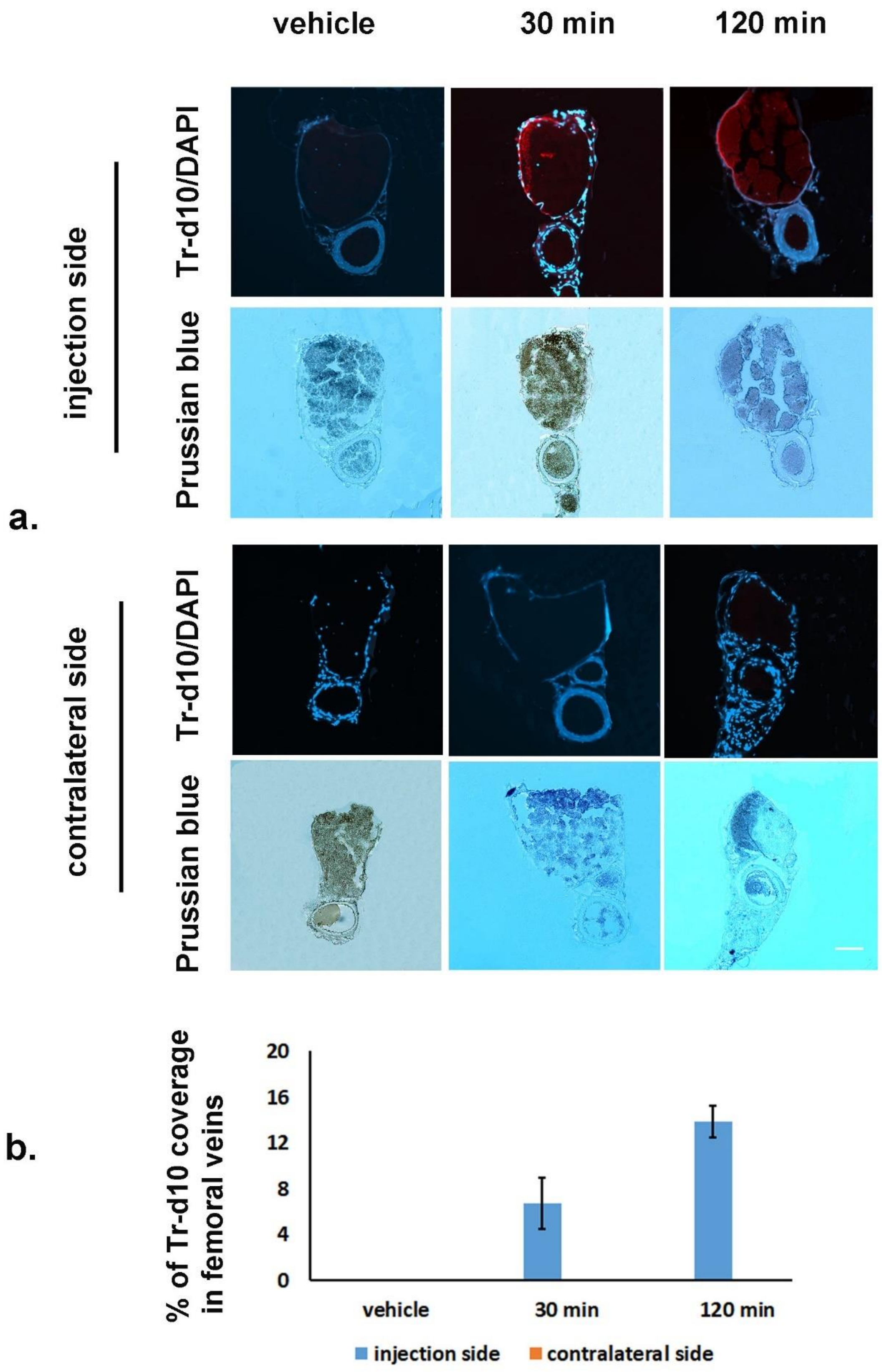

vehicle
30 min
120 min
injection side
contralateral side
Tr-d10/DAPI
Prussian blue
Tr-d10/DAPI
Prussian blue
a.
b.
% of Tr-d10 coverage in femoral veins
20
16
12
8
4
0
vehicle
30 min
120 min
injection side
contralateral side

**Suppl. 8**

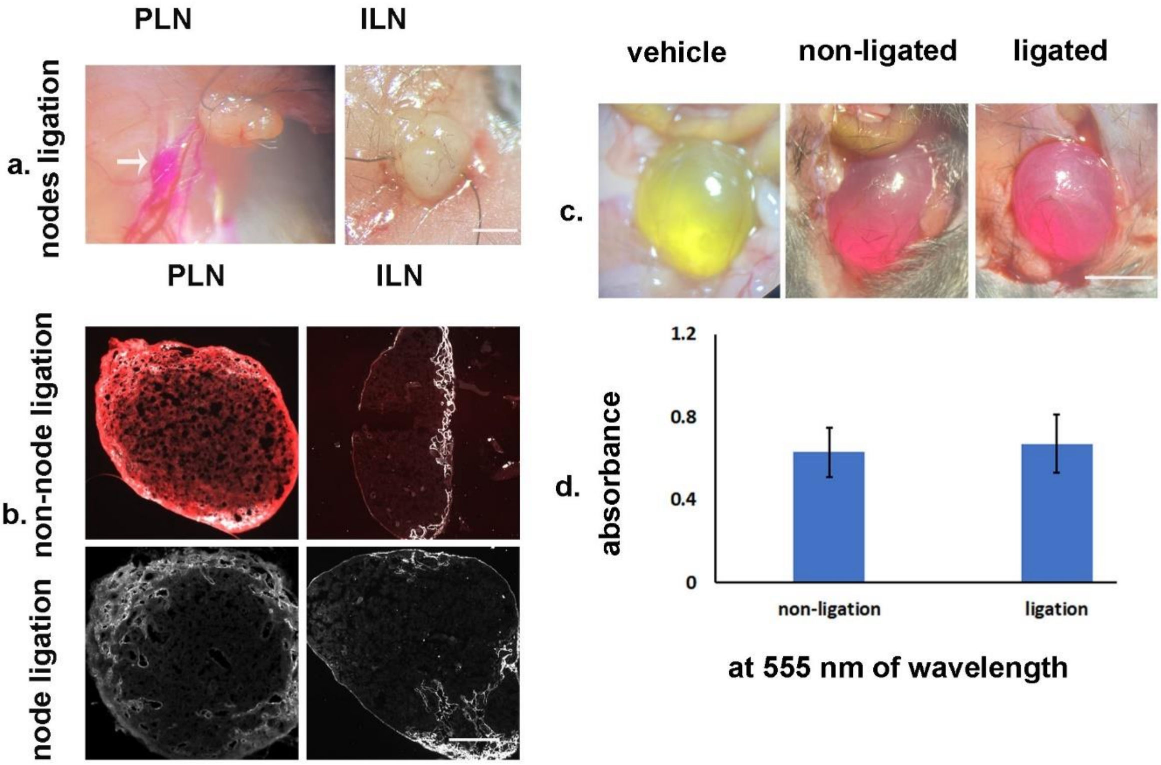


### Supplementary Figure Legends

**Suppl. 1**: ***In Vivo Visualization of CSF Tracer in Prox1 tdTom Mice***

Representative images showing sCLNs and dCLN with their afferent and efferent vessels after ICM injection of OVA488 in Prox1 tdTom mice. The tracer OVA488 is observed within the cervical lymphatic structures of the mice (stars). No tracer is observed in the facial veins and jugular veins. Scale bar represents 1 mm.
